# DNA Damage Response Deficiency Enhances Neuroblastoma Progression and Sensitivity to Combination PARP and ATR Inhibition

**DOI:** 10.1101/2024.09.09.612065

**Authors:** Madeline N. Hayes, Sarah Cohen-Gogo, Lynn Kee, Alex Weiss, Mehdi Layeghifard, Yagnesh Ladumor, Ivette Valencia-Sama, Anisha Rajaselvam, David R. Kaplan, Anita Villani, Adam Shlien, Daniel A. Morgenstern, Meredith S. Irwin

**Author notes:** Correspondence: Madeline N. Hayes, The Hospital for Sick Children PGCRL Building 686 Bay Street, Toronto, Ontario, M5G 0A4, Canada; Meredith S. Irwin, The Hospital for Sick Children PGCRL Building 686 Bay Street, Toronto, Ontario, M5G 0A4, Canada. **Disclosure of Potential Conflicts of Interest:** The authors declare no potential conflicts of interest.

## Abstract

Next generation sequencing of neuroblastoma (NB) tumors have revealed frequent somatic and germline genetic alterations in genes encoding proteins involved in DNA damage response (DDR) pathways. Despite being well-studied in many adult cancers, roles for DDR disruption in pediatric solid tumors remains poorly understood. To address this, patient-relevant loss-of-function mutations in DDR pathway components including Brca2, Atm, and Palb2 were incorporated into an established zebrafish MYCN transgenic model (Tg(*dbh*:EGFP-MYCN)). These mutations were found to enhance NB formation and metastasis *in vivo*, and result in upregulation of proliferation, cell cycle checkpoint and DNA damage repair transcriptional signatures, revealing potential molecular vulnerabilities in DDR-deficient NB. Zebrafish DDR-deficient NB and human NB cells with DDR protein knock-down were sensitive to the poly(ADP-ribose)-polymerase (PARP) inhibitor olaparib, and this effect was further enhanced by inhibition of the ataxia telangiectasia and rad3 related (ATR) kinase. Altogether, our data supports a functional role for DDR-deficiency in NB *in vivo* and therapeutic potential for combination PARP + ATR inhibition in NB patients with alterations in DDR genes.

**Significance:** This work provides the first *in vivo* evidence supporting a functional role for DDR-deficiency in NB by demonstrating that alterations in certain DDR pathway genes promote NB formation and metastasis. NGS and pre-clinical drug testing also provides rationale for PARP + ATR inhibitor therapy combinations for patients with NB and pathogenic DDR pathway alterations.

## Introduction

Neuroblastoma (NB), the most common extra-cranial solid tumor in children, develops from embryonal neural crest-derived precursor cells of the parasympathetic nervous system^1,2^. NB clinical presentations are highly heterogeneous, with outcomes varying from spontaneous tumor regression to aggressive growth and dissemination to multiple sites including bone and bone marrow^3–5^. Despite intensive multi-modal therapies including induction chemotherapy, surgery, high-dose chemotherapy with autologous hematopoietic transplantation, radiation therapy, and immunotherapy, more than half of patients with high-risk disease eventually relapse, most commonly with metastatic disease, and long-term cures following recurrence are rare^1^. Significant work has gone into uncovering the genetic basis of NB, with multiple genetic loci now implicated in germline predisposition, as well as sporadic aggressive disease^6–12^. However, with the exception of the anaplastic lymphoma kinase (*ALK*) gene, which is mutated or amplified in 10-14% of tumors, genomic characterizations of newly diagnosed and relapsed NB have identified very few targetable recurrent alterations^13–17^. Numerous potentially pathogenic/likely pathogenic (P/LP) germline and oncogenic somatic genetic variants have been identified in patients, yet the functional roles for many of these gene mutations in NB pathogenesis and their potential for therapeutic targeting remain poorly understood.

Amplification of the *MYCN* oncogene (*MYCNA*) is detected in approximately 20% of NB tumors and is one the strongest independent prognostic markers of unfavorable outcome^1,18^. The frequency of *MYCNA* is over 40% in high-risk NB tumors; however, even those with *MYCNA* NB display diverse clinical responses, suggesting important underlying biological modifiers affecting tumor growth, invasion, and metastasis^18^. Our previous work identified specific gene expression profiles associated with high-risk *MYCNA* NB, including disruption of DNA damage response (DDR) signaling and upregulation of pathway targets including poly-ADP ribose polymerase 1 (PARP1)^19^. Furthermore, recent next generation sequencing identified multiple somatic and germline alterations in genes encoding essential DDR pathway proteins in pediatric patients enrolled in the SickKids’ Cancer Sequencing (KiCS) program^20^, suggesting that specific genetic events may underly DDR pathway disruption in certain patients with NB. Variants in DDR pathway genes have been described in multiple pediatric oncology patient cohorts, including in CNS tumors, sarcomas, and NB^20–25;^ however, the functional consequence of DDR pathway disruption on NB onset and progression remains to be fully elucidated.

DDR is critical for the maintenance of genomic stability and defects contribute to cancer progression in many adult cancers including breast, ovarian, pancreatic, and prostate^26,27^. Importantly, DDR-deficiency has been shown to increase cellular vulnerabilities to DNA damage and dependencies on remaining repair mechanisms^28,29^. As a result, targeting specific DDR pathways including poly (ADP-ribose) polymerase (PARP) proteins in tumors harboring DDR defects leads to synthetic lethality, offering an effective therapeutic strategy for certain cancers^30–32^. Multiple *in vitro* and *in vivo* studies using human NB cell lines suggests a therapeutic potential for PARP inhibitors (PARPi) in the treatment of high-risk NB patients with DDR pathway alterations, including *ATM, BARD1,* and *ATRX* mutations^33–37^. However, despite important implications for DDR pathway mutations and therapies in adult cancers, the underlying molecular mechanisms supporting DDR-deficiency, impacts on genetic counseling, and targeting DDR-deficiency therapeutically in NB and other pediatric embryonal tumors requires further studies.

Zebrafish are a powerful model system for better understanding biological mechanisms associated with genetic alterations identified in human cancers. An established zebrafish transgenic model of MYCN-driven NB has been impactful with respect to defining roles for candidate genetic modifiers including *ALK*, *LMO1*, and *PTPN11*, among other genes, which are now widely considered important molecular drivers of high-risk NB^38–40^. Given the utility of the MYCN*-*driven zebrafish model and its potential for *in vivo* preclinical drug testing applications, we developed novel engineered zebrafish genetic models, to show that patient-relevant alterations in genes involved in DDR drive unique molecular profiles and offer potentially new therapeutic opportunities for patients with high-risk NB. MYCN-driven zebrafish NB with loss-of-function mutations in *brca2, atm,* and *palb2* display enhanced tumor formation, widespread NB dissemination and metastatic growth. NB tumors from DDR-deficient zebrafish display increased expression of genes associated with cell proliferation, cell cycle checkpoint, and DNA damage repair, and are more sensitive to combination therapies involving the PARPi olaparib. DDR gene knockdown in human NB cells also increases PARPi sensitivity, which is further enhanced by combination treatment with the ATR inhibitor (ATRi) ceralasertib. Altogether, our patient-specific modeling approach supports an important role for DDR pathway mutations in NB and provides rationale for novel therapeutic combinations in the treatment of patients with high-risk NB with DDR pathway alterations.

## Results

### Germline and somatic loss-of-function alterations in DDR pathways are detected in NB tumors

Multiple alterations in DDR have been identified in human NB^19–23,37,41–44^. Notably, recent next generation sequencing of tumor and germline samples from 300 patients (348 tumor samples) enrolled in the SickKids’ Cancer Sequencing (KiCS) Program revealed a high prevalence of alterations in DDR pathway genes across different subtypes of pediatric cancer^20^. This sequencing cohort included 42 NB patients, with ∼24% (10/42) displaying alterations in one or more DDR component^20^. 5/42 (∼12%) had detectable germline variants, with mutations in *BLM, BRCA1, CHEK2, PALB2,* and *BARD1* identified^20^. At the somatic level, mutations were identified in *ATRX, BRCA2, ATM,* and *CHEK1* among other genes, and 14/16 tumor samples had somatic copy number loss of one or more DDR genes (Supplementary Table S1)^20^. Pathogenic/likely pathogenic (P/LP) germline variants were defined based on American College of Medical Genetics and Genomics (ACMG) recommendations, with supporting evidence predominantly from adult cancer studies^45,46^. Loss of heterozygozity (LOH) at *PALB2* was identified in one case with a germline pathogenic variant, and two somatic hits at *BRCA2* in another (Supplementary Table S1); however, LOH was not commonly detected across patients with DDR gene mutations^20^. Importantly, the presence of relevant mutation signatures in NB samples (single-substitution signature 3, SBS3; BRCAness mutational signature), provides additional support for DDR-deficiency in mediating oncogenic processes through mutagenesis (Supplementary Table S1)^20^. However, the functional consequence of these DDR pathway alterations in NB pathogenesis and whether they identify targetable vulnerabilities or increased cancer predisposition risk in patients with NB requires further studies.

### Zebrafish DDR-deficiency increases the penetrance of MYCN-induced neuroblastoma *in vivo*

High genetic conservation and ease of manipulation have made zebrafish a powerful model for studying cancer progression *in vivo*. Transgenic over-expression of MYCN in the peripheral nervous system under the control of the zebrafish *dopamine β-hydroxylase (dbh)* promoter (Tg(*dbh*:EGFP-MYCN)) was previously shown to fluorescently label tumor cells that closely resemble human NB^38^. Therefore, to investigate the role for DDR deficiency in the pathogenesis of NB *in vivo*, we first used a CRISPR-Cas9-mediated gene knockout strategy to disrupt zebrafish homologs of DDR genes altered in patients with NB, in Tg(*dbh*:EGFP-MYCN)^20,47^. Given that disruption of Tp53 is often required for tumorigenesis in the context of DDR deficiency in other models, and that the Tp53 signaling pathway is frequently altered in patients with high-risk NB at relapse, we also incorporated an established loss-of-function *tp53^del/del^* mutant allele (*MYCN;tp53^-/-^*)^48–52^. Our prediction was that a Tp53 loss-of-function background would more robustly reveal the effects of MYCN/DDR-deficiency through inhibition of tp53-mediated cell death activated by MYCN and increased DNA damage resulting from engineered DDR deficiency *in vivo*^53,54^. Furthermore, to enhance the fluorescent intensity of Tg(*dbh*:EGFP-MYCN) and visualization of NB development, we incorporated Tg(*dbh*:EGFP) (*EGFP;MYCN;tp53^-/-^*), as previously described^55^.

We designed 1-3 unique guide RNAs (sgRNAs) targeting coding sequences of *brca2, atm, palb2,* and *bard1* and co-injected sgRNAs with Cas9 protein into *EGFP;MYCN*;*tp53^-l-^* embryos, at the one-cell stage (Supplementary Fig 1A, Supplementary Table S2). Prior to co-injection, each gRNA was independently verified and displayed an indel rate of >90%, suggesting efficient gene targeting (data not shown, ICE Analysis, Synthego Performance Analysis). Starting at 6 weeks post fertilization (wpf), mosaic primary injected and un-injected control *EGFP;MYCN;tp53^-/-^* animals were screened for EGFP-positive lesions at the site of the interrenal gland (Supplementary Fig 1B-E), equivalent to the adrenal gland in humans^38,56^. *EGFP;MYCN;tp53^-/-^* control fish developed NB with an overall incidence of 20-30% (Supplementary Fig 1F). In contrast, mosaic primary *brca2, atm, palb2,* and *bard1* CRISPR/Cas9-injected animals displayed increased EGFP-labeled lesions, with up to 63% of animals developing NB by 22 weeks of age (Supplementary Fig 1F, Log-rank test). Randomly selected tumor-burdened mosaic injected fish from each cohort were sacrificed at 12-15 weeks of age and EGFP+ tumor cells isolated using fluorescent activated cell sorting (FACS). Sequencing was used surrounding gRNA target sites and mutations predicted to affect protein function through large deletions and/or frameshift mutations were found in each of *brca2, atm, palb2,* and *bard1* (Supplementary Fig 1B-E, Supplementary Table 3), suggesting enhanced NB formation following DDR gene targeting *in vivo*.

To further examine the role for DDR deficiency in NB and validate the effects of DDR gene targeting *in vivo*, we next established mutant zebrafish lines with germline transmission of predicted loss-of-function alleles in *brca2, atm, palb2*. From outcrossing our mosaic primary CRISPR/Cas9-injected cohorts to un-injected *tp53^+/-^* or *tp53^-/-^* animals, we recovered a large deletion at the *brca2* locus upstream of exon 1 and the predicted translational start site in exon 2, through to exon 10 (Supplementary Fig 2A Supplementary Table 2). For *atm*, we recovered a four base pair deletion in exon 54, resulting in a frame shift mutation and pre-mature stop codon upstream of the kinase domain (Supplementary Fig 2A, Supplementary Table 2). For *palb2*, we recovered a large deletion between exon 2 and exon 5, which deleted the translational start site (Supplementary Fig 2A, Supplementary Table 2). Previously reported zebrafish strains carrying mutations in individual DDR genes, including *brca2, atm* and *palb2*, are strongly biased towards the male sex due to an important role for these genes in meiotic recombination, oocyte viability, differentiation, and sex determination^57–61^. Tp53 loss was shown to rescue germ cell apoptosis and male sex bias, albeit with later-stage germ-cell tumor formation in *brca2* mutants^60,61^. Consistent with these previous reports, we found that our *brca2, atm* and *palb2* homozygous mutant zebrafish strains were 100% males in *tp53* wild-type and heterozygous backgrounds, and that *tp53^-/-^* loss could variably rescue this effect (Supplementary Fig 2B). *Brca2* and *atm* homozygous males were also sterile when bred to heterozygous or wild-type zebrafish. *Brca2* and *palb2* alleles were recovered at Mendelian ratios up to 4 months of age, while *atm^-/-^;tp53^-/-^* zebrafish displayed early mortality starting at 1 month post fertilization (Supplementary Fig 2C), consistent with early mortality potentially linked to a developmental blockade in hematopoiesis and associated immunological defects^57^. Furthermore, we found that *brca2^-/-^*;*tp53^-/-^* females were partially fertile and bred rare viable embryos with *brca2^+/-^* zebrafish that displayed severe developmental abnormalities and embryonic lethality prior to 24 hours post fertilization (24hpf) (Supplementary Fig 2D), consistent with an important role for DDR proteins in vertebrate germ cells and development across model systems^50,61,62^. Altogether, these observations suggest strong loss-of-function alleles in our germline zebrafish models and important roles for DDR proteins throughout development and juvenile stages.

To determine whether loss of Brca2 and/or other DDR genes affects tumor development we utilized MYCN transgenic zebrafish NB with genetically engineered alterations in DDR pathway genes. After interbreeding *EGFP;MYCN;tp53^-/-^;brca2^+/-^* to *tp53^-/-^;brca2^+/-^* adult zebrafish we screened progeny for EGFP+ tumor development starting at 6 weeks up to 22 weeks of age. Notably, 49% of *brca2* heterozygous (*EGFP;MYCN;tp53^-/-^;brca2^+/-^*) and 85% of *brca2* homozygous (*EGFP;MYCN*;*tp53^-/-^*;*brca2^-/-^*) mutant animals developed EGFP+ tumors, compared with an overall penetrance of 26% for *brca2* wild-type (*EGFP;MYCN;tp53^-/-^*) alone (Fig 1A,B; *brca2^+/-^* p=0.0289 and *brca2^-/-^* p<0.0001, Log-rank test). *Tp53* loss-of-function was required for increased tumor formation observed with *brca2* mutation, prior to 22 weeks of age (Supplementary Fig 3), supporting a role for combined cell cycle disruption and DDR-deficiency in enhancement of tumorigenesis *in vivo*. A similar breeding strategy was used to assess *atm* and *palb2* mutant alleles, with 76% of *atm* heterozygous (*EGFP;MYCN;tp53^-/-^;atm^+/-^*) and 100% of *atm* homozygous (*MYCN;tp53^-/-^;atm^-/-^*) mutant animals developing EGFP+ tumors (Fig 1C, *atm^+/-^* p=0.0004 and *atm^-/-^* p<0.0001, Log-rank test), and 60% of *palb2* homozygous mutant (*EGFP;MYCN;tp53^-/-^;palb2^-/-^*) developing EGFP+ tumors prior to 22 weeks (Fig 1D, p=0.0222, Log-rank test).

**Figure 1.**
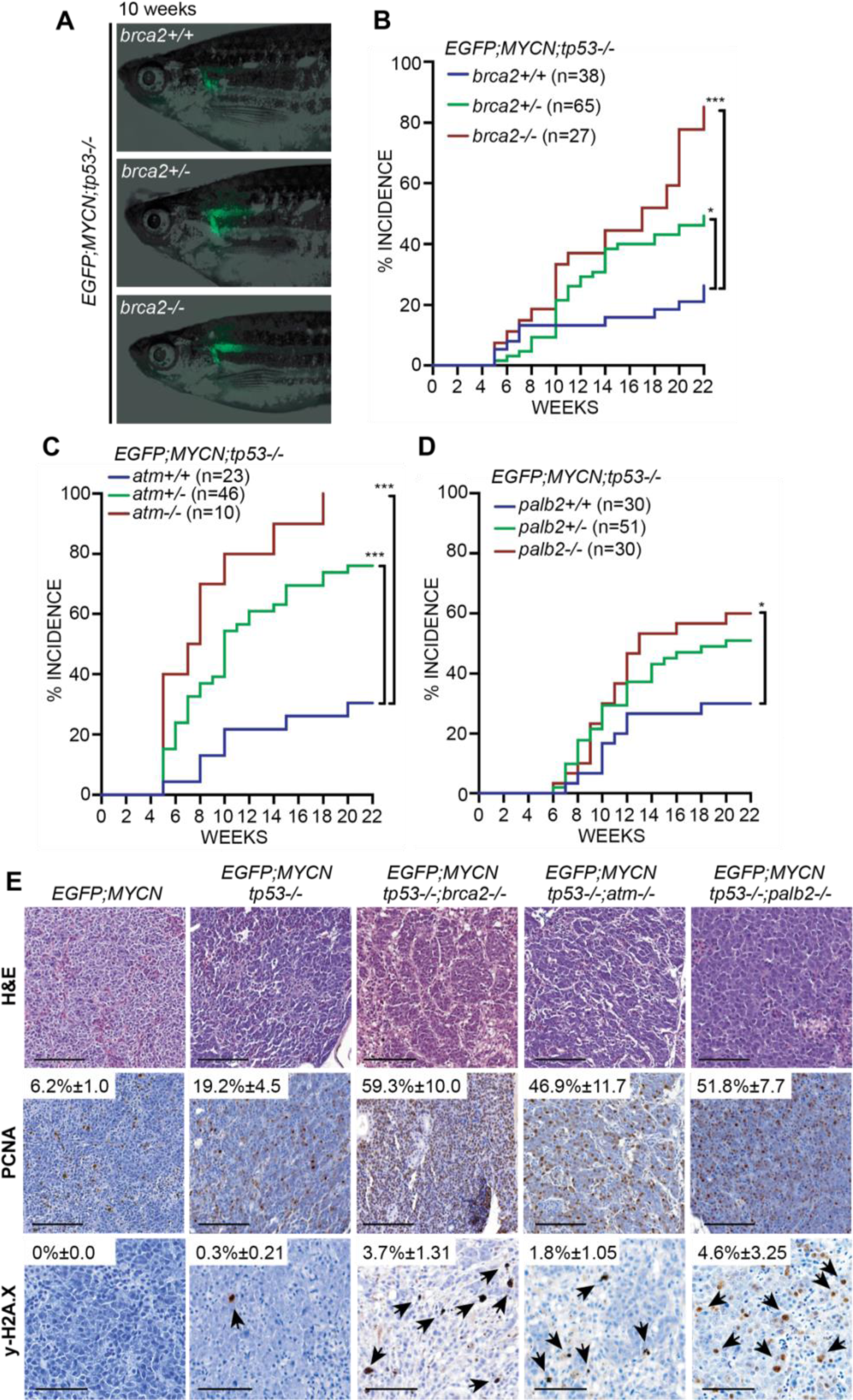
Inherited germline DDR-deficiency enhances MYCN-induced NB formation. **(A)** EGFP expression in primary NB localized at the interrenal gland of 10 week post fertilization (wpf) *EGFP;MYCN;tp53^-/-^;brca2^+/+^, EGFP;MYCN;tp53^-/-^;brca2^+/-^,* and *EGFP;MYCN;tp53^-/-^;brca2^-/-^* zebrafish. **(B-D)** Cumulative frequency of NB onset among *EGFP;MYCN;tp53^-/-^* cohorts carrying wild-type and mutant alleles of *brca2* (B), *atm* (C), and *palb2* (D). Differences between curves are compared to *EGFP;MYCN;tp53^-/-^* wild-type siblings. *p<0.05, ***p<0.001, Log-rank test. **(E)** Hematoxylin and eosin (H&E) staining and immunohistochemical staining of PCNA and y-H2A.X of sections from primary NB of *EGFP;MYCN, EGFP;MYCN;tp53^-/-^, EGFP;MYCN;tp53^-/-^;brca2^-/-^, EGFP;MYCN;tp53^-/-^;atm^-/-^,* and *EGFP;MYCN;tp53^-/-^;palb2^-/-^* animals. Quantification of positive PCNA nuclei relative to total is indicated and is increased compared to *EGFP;MYCN* and *EGFP;MYCN;tp53^-/-^* (*brca2^-/-^*p=0.0032, *atm^-/-^* p=0.018, *palb2^-/-^* p=0.0032, n=3). Quantification of positive y-H2A.X nuclei relative to total is indicated (*brca2^-/-^* p=0.0081, *atm^-/-^* p=0.0412, *palb2^-/-^* p=0.0703, compared to *EGFP;MYCN*, n=3). Black arrowheads indicate y-H2A.X-positive nuclei. Scale bars = 50μm.

*EGFP;MYCN, EGFP;MYCN;tp53^-/-^*, and *EGFP;MYCN;tp53^-/-^*;DDR-deficient tumors arose at the site of the interrenal gland and were composed of undifferentiated, small, round blue tumor cells with hyperchromatic nuclei, comparable with human NB tumors and previously described MYCN-driven animal models of NB (Fig 1E)^38,63,64^. NB tumors with gene mutations in DDR genes displayed increased proportions of PCNA-positive nuclei, compared to *EGFP;MYCN* and *EGFP;MYCN;tp53^-/-^* NB (Fig 1E), suggesting increased proliferation in DDR-deficient backgrounds. In contrast to *EGFP;MYCN* tumor sections, at 3-5 months we observed immunohistological staining of phosphorylated Histone H2A.X (S139, y-H2A.X), a marker of double stranded DNA damage, in *EGFP;MYCN;tp53^-/-^* and *EGFP;MYCN;tp53^-/-^;*DDR*-*deficient models (Fig 1E), suggesting increased DNA damage and/or diminished repair mechanisms. Altogether, these findings suggest that patient-relevant loss of DDR genes leads to reduced DSB break repair and highly proliferative sympathoadrenal-derived tumor cells *in vivo*, contributing to the increase in penetrance of MYCN-driven NB in zebrafish.

### Zebrafish DDR-deficiency promotes metastasis of MYCN-induced neuroblastoma *in vivo*

Alterations in DDR signaling pathway genes were identified in our NB patient cohort, which included a majority of patients with relapsed or refractory high-risk disease, who more often display metastatic disease^20^. To assess whether DDR-deficiency directly promotes metastasis *in vivo*, we screened control and *EGFP;MYCN;tp53^-/-^*;DDR-deficient zebrafish for metastases, using disseminated EGFP+ masses distant from primary tumors as a general indicator of metastatic burden. Notably, and in contrast to DDR competent models, we detected frequent and wide-spread NB dissemination among tumor-bearing *EGFP;MYCN;tp53^-/-^;*DDR-deficient zebrafish carrying both heterozygous and homozygous germline mutations in *brca2*, *atm*, and *palb2* prior to 22 weeks of age (Fig 2A,B; Supplementary Fig 4A). DDR-deficient animals displayed a significant increase in distant lesions both anterior and posterior to the interrenal gland compared to *EGFP;MYCN;tp53^-/-^*, with up to 100% of tumor-bearing *EGFP;MYCN;tp53^-/-^;*DDR-mutant animals displaying widespread EGFP-positive tumor cells at required end-point (Fig 2B, *brca2^+/-^* p=0.031, *brca2^-/-^* p<0.0001, *atm^+/-^* p<0.0001, *atm^-/-^* p<0.0001, *palb2^+/-^* p=0.0154, *plab2^-/-^* p=0.0235, Fisher exact test). Consistent with previous reports, *EGFP;MYCN* zebrafish NB did not display detectable distant EGFP+ metastases prior to 1 year of age (Fig 2B)^65^, and we detected only the occasional occurrences of disseminated EGFP in *EGFP;MYCN*;*tp53^-/-^* animals prior to 22 weeks of age, visualized by fluorescent cell masses localized mainly to the interrenal gland (Fig 2B, Supplementary Fig 4B).

**Figure 2.**
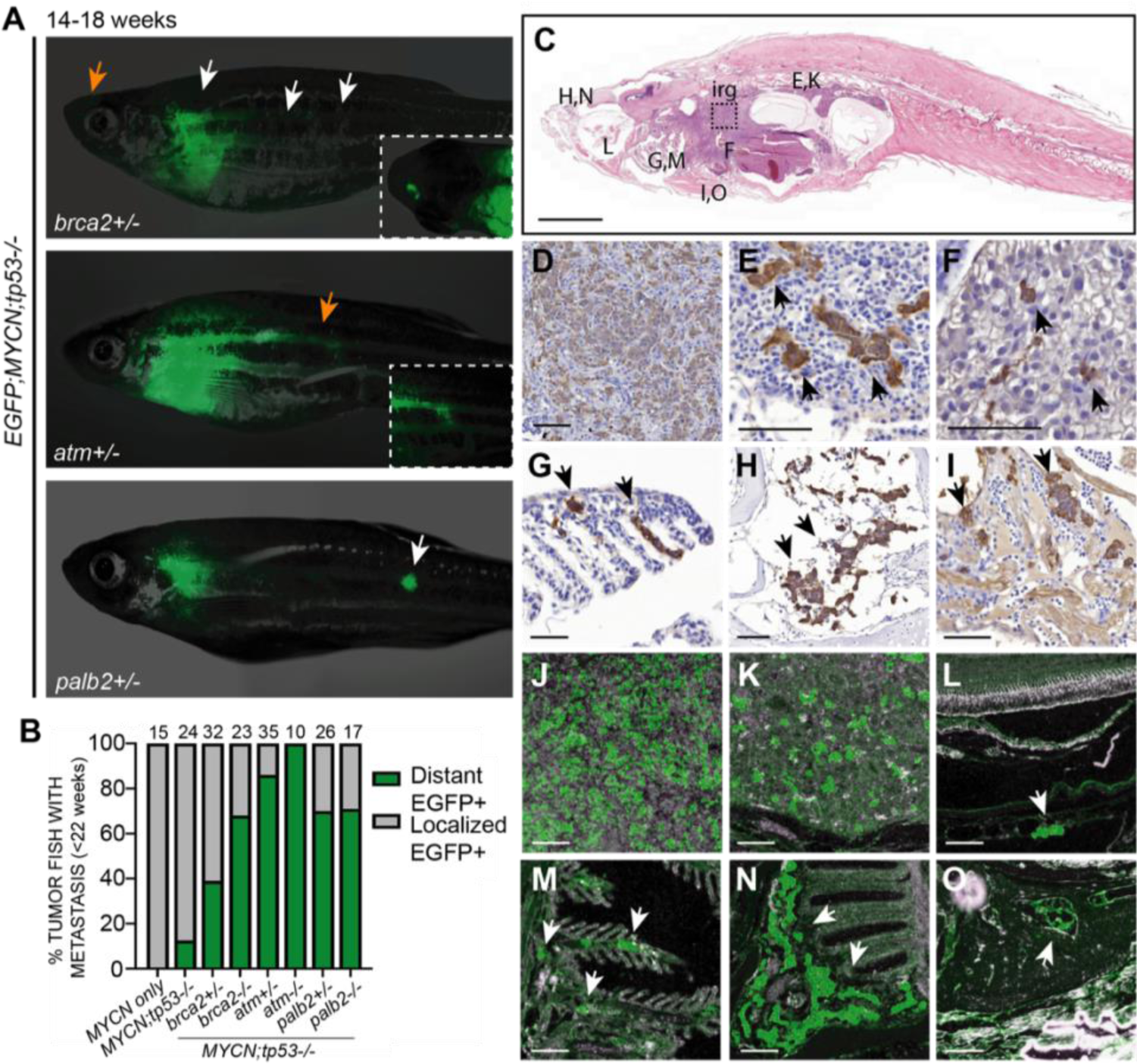
DDR-deficiency promotes NB metastasis *in vivo*. **(A)** EGFP fluorescence images of *EGFP;MYCN;tp53^-/-^;brca2^+/-^, EGFP;MYCN;tp53^-/-^;atm^+/-^,* and *EGFP;MYCN;tp53^-/-^;palb2^+/-^* zebrafish at 14-18 wpf displaying disseminated lesions (arrows). Insets highlight regions indicated by orange arrows. **(B)** Proportion of tumor-bearing zebrafish displaying metastases prior to 22 weeks of age. The percentage of DDR-deficient animals with disseminated EGFP is significantly increased compared to *EGFP;MYCN;tp53^-/-^* by Fisher exact test (*brca2^+/-^* p=0.031, *brca2^-/-^* p<0.0001, *atm^+/-^* p<0.0001, *atm^-/-^* p<0.0001, *palb2^+/-^* p=0.0154, *plab2^-/-^* p=0.0235). **(C)** Hematoxylin and eosin (H&E)-stained sagittal section of a representative *EGFP;MYCN;tp53^-/-^;brca2^-/-^* at 20 weeks of age. **(D-I)** Immunohistological stains of sagittal sections of *EGFP;MYCN;tp53^-/-^;brca2^-/-^* using tyrosine hydroxylase (TH). **(J-O)** Immunofluorescence stains of *EGFP;MYCN;tp53^-/-^;brca2^-/-^* using anti-GFP. Black dotted box in (C) outlines the interrenal gland (irg) that is stained and magnified in (D) and (J). Renal tubules are stained and magnified in (E) and (K). Disseminated tumor cells were also detected in DDR-deficient models in the liver (F, arrow), the orbit (L, arrow), the gills (G, M, arrows), olfactory pits (H, N, arrows) and within the heart chambers (I, O, arrows). Scale bars = 100μm.

To validate and further investigate the organ-specific locations of disseminated NB cells in our models, we performed sectioning on tumor-bearing *EGFP;MYCN;tp53^-/-^* and *EGFP;MYCN;tp53^-/-^;brca2^-/-^* zebrafish (Fig 2C), and visualized tumor cells using immunohistological staining of tyrosine hydroxylase (TH) (Fig 2D-I), a common marker of human NB, and GFP (Fig 2J-O), indicating cells expressing Tg(*dbh:*EGFP) and Tg(*dbh:*EGFP-MYCN). Consistent with previous descriptions of MYCN-driven zebrafish NB^39^, multifocal nests of TH and GFP-positive tumor cells were detected in the interrenal glands and renal tubules of *EGFP;MYCN;tp53^-/-^* and mutant *EGFP;MYCN;tp53^-/-^;brca2^-/-^* (Fig 2C-E, 2J-K and Supplementary Fig 4B). In *EGFP;MYCN;tp53^-/-^;brca2^-/-^* animals, we also observed TH and GFP-positive NB lesions in the liver, orbits, gills, and olfactory pits (Fig 2C, 2F-H, 2L-N, Supplementary Fig 4B), supporting widespread and multi-organ metastasis. We also observed positive staining within the inner wall of heart chambers, suggesting hematogenous NB dissemination (Fig 2I,O; Supplementary Fig 4B). Therefore, in addition to enhanced NB formation, enhanced metastatic behavior in DDR-deficient zebrafish supports a role for DDR in NB progression, significantly in hematogenous dissemination and growth at distant secondary organs, which in patients is a major predictor of poor clinical outcomes.

### PARP inhibition suppresses DDR-deficient NB growth

Olaparib is an orally bioavailable inhibitor of PARP1 and PARP2, which have key roles in single-stranded DDR^66,67^. Blocking PARP activity shifts the reliance of cells to repair DNA breaks using homology directed repair (HDR). In tumors also lacking BRCA1, BRCA2, PALB2 or ATM, there is inefficient repair of DNA breaks, leading to cell death and inhibition of tumor growth^28,68–70^. Additionally, PARP inhibitors have been shown to be more effective in combination with DNA-damaging chemotherapies. To determine whether the PARP inhibitor olaparib, when combined with TMZ chemotherapy, is effective at inhibiting DDR-deficient NB growth *in vivo*, we dosed single agent and combination drugs to *EGFP;MYCN;tp53^-/-^;brca2^-/-^* homozygous mutant DDR*-*deficient NB cells.

To generate sufficient zebrafish with tumors for comparative drug trials we expanded primary *EGFP;MYCN;tp53^-/-^;brca2^-/-^* NB using allograft transplantation of primary zebrafish NB cells into 15-30 immune-deficient *casper;prkdc^-/-^* hosts^71^. Engrafted animals were selected 7-14 days post transplantation using epifluorescence and pre-treatment tumor sizes were quantified using EGFP+ NB area multiplied by fluorescent intensity, as previously described (Fig 3A)^72^. Engrafted animals were then orally gavaged 4 consecutive days per week (1 dose per day, 4 days on, 3 days off), for a total of 21 days with either DMSO vehicle control, olaparib (50mg/kg), TMZ (33mg/kg), or both drugs (50mg/kg olaparib + 33mg/kg TMZ). Oral gavage of TMZ + olaparib in zebrafish was previously shown to result in plasma concentrations similar to humans and preclinical mouse models, as well as effectively inhibit tumor growth of other sensitive cancer subtypes *in vivo*^72^. Control DMSO-treated animals displayed robust EGFP+ NB growth, and single agent treatments resulted in moderate inhibition of tumor growth *in vivo* (Fig 3A-C). Notably, olaparib + TMZ combination treatment led to significant inhibition of *EGFP;MYCN;tp53^-/-^;brca2^-/-^* tumors, compared to DMSO vehicle control and single agent olaparib or TMZ treatments (Fig 3A,B; *brca2^-/-^* p=0.0016 DMSO vs. olaparib + TMZ, n=3 independent primary tumors). In contrast, at the same doses olaparib + TMZ combination treatment of fish bearing *brca2* wild-type (*EGFP;MYCN;tp53^-/-^*) NB cells had an insignificant effect on tumor growth, compared to DMSO vehicle-control treated animals (Fig 3C, n=2 independent primary tumors). Histological evaluations confirmed overall reductions in cellularity on sections in animals treated with olaparib + TMZ, as well as proliferative cells marked by PCNA, in all treatment groups (Fig 3D, Supplementary Fig 5). Increased cleaved caspase-3 (CC3) positive cells were detected in combination olaparib + TMZ treated tumors (Fig 3D), suggesting on-target drug-induced cytotoxicity in *brca2* homozygous loss-of-function DDR mutant NB *in vivo*.

**Figure 3.**
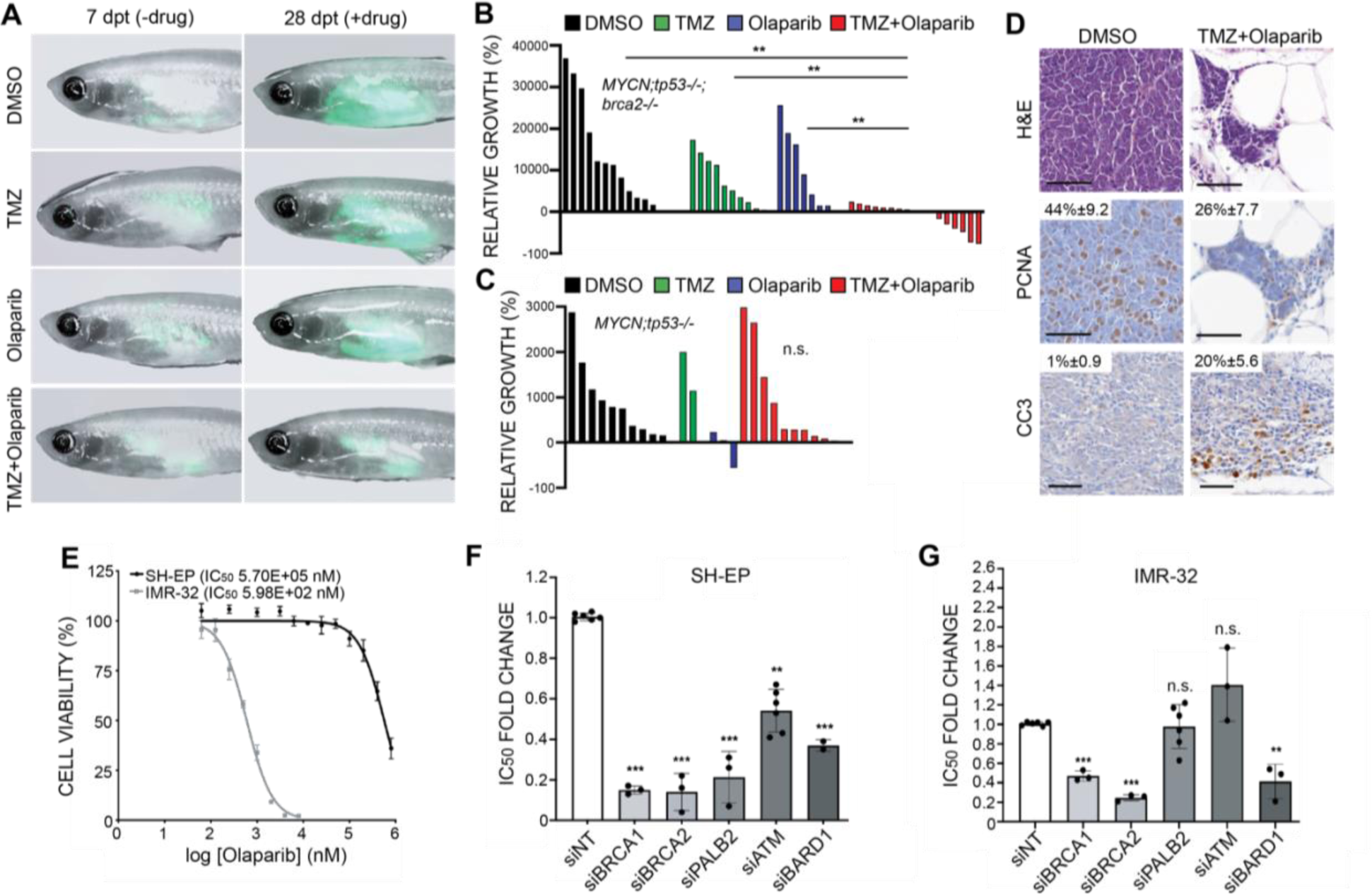
DDR-Deficiency enhances NB sensitivities to Olaparib PARP Inhibitor with and without Temozolomide (TMZ) Chemotherapy. **(A)** *casper^-/-^;prkdc^-/-^* animals engrafted with *EGFP;MYCN;tp53^-/-^;brca2^-/-^* NB cells at 7 days post transplant (dpt) prior to oral drug administrations, and at 28 dpt after three doses of DMSO vehicle control, TMZ, olaparib, or olaparib + TMZ combination therapy. **(B,C)** Quantification of relative growth after three doses (28 dpt) of DMSO vehicle control, TMZ, olaparib, or olaparib + TMZ to *EGFP;MYCN;tp53^-/-^;brca2^-/-^* NB ((B), n=3 independent primary tumors), and *EGFP;MYCN;tp53^-/-^* NB ((C), n=2 independent primary tumors). **p<0.01, not significant (n.s.), Student’s t-test. **(D)** Hematoxylin and eosin (H&E), PCNA, and Cleaved caspase 3 (CC3) staining on *EGFP;MYCN;tp53^-/-^;brca2^-/-^* NB sections following treatment with DMSO vehicle control and combination olaparib + TMZ. The average percent positive cells with standard deviation are noted. Significant CC3 increases were observed in olaparib +TMZ treated *EGFP;MYCN;tp53^-/-^;brca2^-/-^* engrafted NB compared to DMSO control (p=0.0044, n=3). **(E)** Cell viability (AlamarBlue) analysis in IMR-32 and SH-EP human NB cell lines following treatment with Olaparib [0.05-800uM] for 72 hours. **(F)** Determination of half-maximal inhibitory concentration (IC_50_) fold change (normalized to siNT control) in SH-EP cells with DDR pathway knock-down and treated with Olaparib for 72 hours. Error bars, mean ± standard deviation. **p<0.01, ***p<0.001. **(G)** Determination of half-maximal inhibitory concentration (IC_50_) fold change (normalized to siNT control) in IMR-32 cells with DDR pathway knock-down treated with Olaparib for 72 hours. Error bars, mean ± standard deviation. **p<0.01, ***p<0.001.

To test whether DDR-deficiency may enhance sensitivity to PARP inhibition in human NB cells, we used siRNA-mediated knock-down of *BRCA1, BRCA2, PALB2, ATM,* and *BARD1*, followed by olaparib treatment to determine half-maximal inhibitory concentrations (IC_50_). Consistent with our previous work demonstrating that cells harboring *MYCN* amplifications were more sensitive to PARP inhibitors, we found that IMR-32 cells (*MYCNA, TP53* wild-type) were sensitive to olaparib treatment *in vitro*^19,33^, while SH-EP cells (non-*MYCNA*, *TP53* wild-type) were relatively resistant (Fig 3E). DDR protein knock-down in SH-EP cells was associated with increased sensitivity to olaparib, as shown by a reduction in IC_50_ values, as compared to siRNA non-targeting control-transfected cells (Fig 3F, Supplementary Fig 6A), suggesting that DDR-deficiency can increase sensitivity to olaparib in otherwise highly resistant NB cells. *BRCA1, BRCA2,* and *BARD1* knock-down further increased olaparib sensitivity in IMR-32 cells; however, no significant effects were observed following *ATM* or *PALB2* knock-down (Fig 3G, Supplementary Fig 6), suggesting pathway-specific effects in certain NB cell lines related to underlying genetic alterations^33^.

### DDR-deficient NB models display genomic instability and upregulation of cell cycle checkpoint gene expression signatures

HDR following DNA double-strand breaks is essential for maintaining genomic integrity. To better understand underlying genetic mechanisms associated with DDR-deficiency in NB and in our models, we first assessed the genome-wide impact of loss-of-function *tp53* and *brca2* alleles on MYCN-driven zebrafish NB compared to matched normal tissue. We dissected tumors and sorted EGFP-fluorescent primary NB cells as well as EGFP-negative non-tumor tissue from one *EGFP;MYCN*, five *brca2* wild-type (*EGFP;MYCN;tp53^-/-^*), three *brca2* heterozygous mutant (*EGFP;MYCN;tp53^-/-^;brca2^+/-^*), and three *brca2* homozygous mutant (*EGFP;MYCN;tp53^-/-^;brca2^-/-^*) zebrafish. We extracted genomic DNA and performed whole-genome sequencing (WGS) followed by assessments for single-nucleotide variants (SNVs), small insertions and/or deletions (INDELs), and larger structural rearrangements including deletions, insertions, duplications, and translocations (Supplementary Fig 4A-C, Supplementary Table 4, Supplementary Table 5). We observed evidence of genomic instability across models with many SNVs, INDELs, and structural rearrangements. However, *EGFP;MYCN;tp53^-/-^;brca2^+/-^* and *EGFP;MYCN;tp53^-/-^;brca2^-/-^* tumors appeared to more often display an excess of SNVs, INDELs and structural rearrangements, compared to *EGFP;MYCN;tp53^-/-^*, suggesting enhanced HDR defects with *brca2* loss (Supplementary Fig 4). Interestingly, *brca2* heterozygous and homozygous mutant tumors displayed more INDELs (Supplementary Fig 4A,B), compared to *MYCN and MYCN;tp53^-/-^* tumors, consistent with the observation that *BRCA1/2* tumors typically display elevated small INDELs (>3bp)^73^. In coding regions, 6/6 *EGFP;MYCN;tp53^-/-^;brca2^+/-^* and *EGFP;MYCN;tp53^-/-^;brca2^-/-^* NBs displayed 3 or more in-frame INDELs and/or frameshift mutations, compared to 2/5 *EGFP;MYCN;tp53^-/-^* tumors, suggesting increased HDR defects and compensatory alternative repair mechanisms (Supplementary Fig 4B). Furthermore, 5/6 *brca2-*deficient tumors displayed 40 or more structural rearrangements, compared to 2/5 *EGFP;MYCN;tp53^-/-^* (Supplementary Fig 4C), altogether suggesting genomic instability in *brca2*-deficient NB, which corresponds to our observation of enhanced NB tumor formation and metastases *in vivo* (Fig 1,2). Increased genomic instability in our DDR-deficient models is also consistent with observations made in NB sequencing studies demonstrating increased tumor mutational burden and mutational signatures associated with BRCA1/2 deficiency (SBS3)^20^.

Given evidence of genomic instability, we next performed gene expression signature analysis following transcriptome profiling by RNA-seq of EGFP+ sorted *brca2* wild-type (*EGFP;MYCN;tp53^-/-^*), *brca2* heterozygous (*EGFP;MYCN;tp53^-/^;brca2^+/-^*), and *atm* heterozygous mutant (*EGFP;MYCN;atm^+/-^*) NB tumor cells to gain further insight into conserved regulatory pathways involved in maintenance and progression of our aggressive DDR-deficient NB models. Heterozygous loss-of-function DDR models were prioritized for further studies given their aggressiveness and the observation of hemizygous loss-of-function mutational status of DDR factors in the majority of patients in our previous sequencing cohort (Supplementary Table 1). We also analyzed sorted EGFP+ tumor cells from adult *prkdc^-/-^* immune-deficient zebrafish engrafted with *EGFP;MYCN;tp53^-/^;brca2^+/-^* NB. Transplanted *EGFP;MYCN;tp53^-/^;brca2^+/-^* NB cells were considered in order assess roles for DDR-deficiency and potential drug sensitivities following engraftment, which is often used as a surrogate of relapsed tumor growth *in vivo*^74,75^. Through comparisons of primary tumors with differing genotypes we detected enrichment for proliferation signatures including E2F targets, G2M checkpoint, and MYC targets in *brca2^+/-^* and *atm*^+/-^ tumor cells, compared to *EGFP;MYCN;tp53^-/-^* control cells (Fig 4A, Supplementary Table 6). Interestingly, we observed further enrichment of E2F targets and G2/M cell cycle gene expression following *brca2-*deficient tumor transplantation (Fig 4A,B; Supplementary Table 6), suggesting selection for these pathways upon relapsed tumor growth. Together, *brca2* and *atm* DDR-deficient primary NB displayed strong upregulation of genes involved in DNA replication, cell cycle checkpoint pathways, DNA repair, stress responses, and signatures previously associated with *BRCA1* loss in human breast cancers, consistent with HDR-deficiency in our models (Fig 4C, Supplementary Table 6). Interestingly, *brca2^+/-^* tumors also displayed strong enrichment for complement, collagen synthesis, and extracellular matrix regulation, suggesting potential molecular drivers of observed metastases (Fig 4D, Supplementary Table 6). Furthermore, in *EGFP;MYCN:tp53^-/-^;brca2^+/-^* NB, we observed up-regulation of the senescence gene *cdkn1a* (p21) as well as interferon signaling pathways (Supplementary Table 6, Supplementary Table 7), suggesting potential survival mechanisms. In contrast, *EGFP;MYCN;p53^-/-^* control tumors enriched for signatures associated with RAC1 activation, myogenesis, RAS/MAPK signaling, and the TP53 signaling pathway (Figure 4C, Supplementary Table 6), with TP53 signatures previously associated in patient tumors with genetic alterations in TP53^76,77^. Altogether, our primary and transplanted *EGFP;MYCN;tp53^-/-^;*DDR-deficient NB models display up-regulation of genes associated with a variety of pro-tumorigenic signatures including genes previously investigated in NB like *CHEK1, RRM2, PRKDC, WEE1, AURKA,* and *EZH2*^78–87^. Altogether, our genomic and transcriptional analyses suggest enhanced DNA damage and repair mechanisms *in vivo*, which could enhance vulnerabilities to DNA repair and/or cell-cycle checkpoint targeted therapeutics.

**Figure 4.**
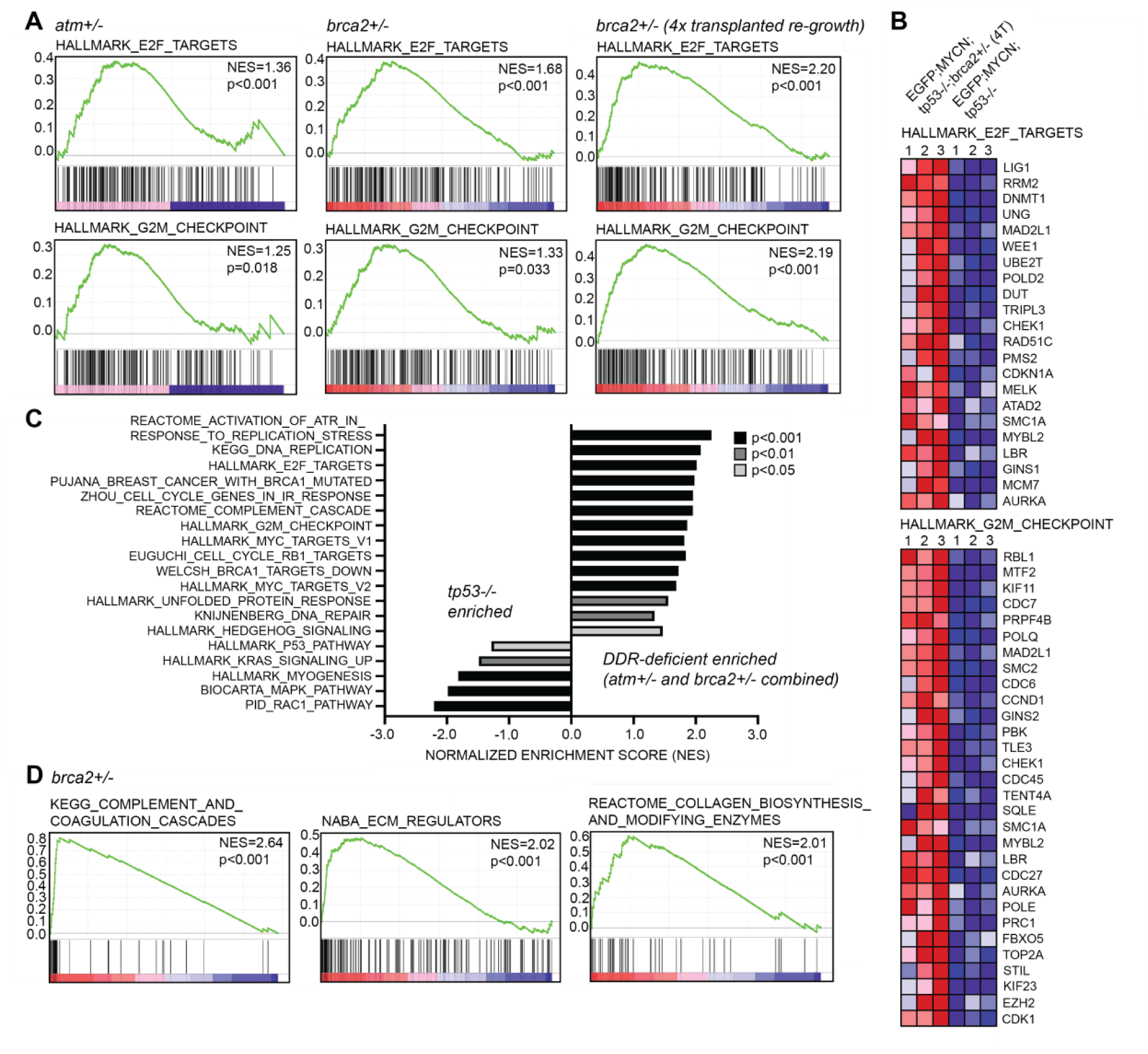
DDR-deficient zebrafish NB tumors display upregulation of DNA repair and cell cycle checkpoint gene expression signatures. **(A)** Gene set enrichment analysis (GSEA) following RNA sequencing-based transcriptome profiling of primary *EGFP;MYCN;tp53^-/-^;atm^+/-^,* primary *EGFP;MYCN;tp53^-/-^;brca2^+/-^,* and four-time (4x) serially transplanted *EGFP;MYCN;tp53^-/-^;brca2^+/-^* NB compared to *EGFP;MYCN;tp53^-/-^*. GSEA indicates enrichment for E2F target genes and G2M checkpoint gene expression signatures. Normalized enrichment scores (NES) and nominal p-values are indicated. **(B)** Heat maps for top enriched E2F and G2M checkpoint genes in transplanted *EGFP;MYCN;tp53^-/-^;brca2^+/-^* NB compared to *EGFP;MYCN;tp53^-/-^*. **(C)** Selected gene signatures enriched in primary NBs. DDR-deficient NB represents *brca2^+/-^* and *atm^+/-^* combined GSEA analysis (5 tumors total). Bar color indicates significance. **(D)** GSEA analysis of primary *EGFP;MYCN;tp53^-/-^;brca2^+/-^* NB compared to *EGFP;MYCN;tp53^-/-^* indicates enrichment for gene expression associated with tumor cell invasion and metastasis. Normalized enrichment scores (NES) and nominal p-values are indicated.

### Targeting the CDK4/6 pathway in combination with PARP inhibition in DDR-deficient NB

Several preclinical modeling studies support inhibitors of cyclin dependent kinase 4/6 (CDK4/6) as treatments for neuroblastoma on the basis of their anti-proliferative and pro-differentiation functions^88–92^. Given upregulation of proliferation signatures, *cyclinD1* and E2F target gene expression in our DDR-deficient zebrafish NB models (Fig 4, Supplementary Table 6, Supplementary Table 7), we decided to investigate the potential for cell cycle suppression through CDK4/6 inhibition as a therapeutic vulnerability further enhanced by DDR gene loss-of-function in NB. Specifically, we were interested in whether palbociclib, an inhibitor of CDK4/6 (Ibrance, Pfizer; PD-0332991) could synergize with olaparib PARP inhibitor in both human NB cells and zebrafish models. We again assessed PARPi-resistant SH-EP cells following siRNA-mediated knock-down of *BRCA1, BRCA2, PALB2, ATM,* and *BARD1* (Supplementary Fig 6). The addition of 100nM palbociclib enhanced olaparib-mediated inhibition of SH-EP growth in *BRCA2* and *PALB2* knock-down cells, as seen by reduction in IC_50_ values compared to olaparib treatment alone and compared to olaparib + palbociclib treatments in control siRNA treated cells (Fig 5A, Supplementary Table 8). In contrast, combination olaparib + palbociclib treatments following *ATM*, *BRCA1*, and *BARD1* gene knock-down in SH-EP cells did not enhance olaparib-mediated inhibition, compared to control siRNA treated cells (Fig 5A, Supplementary Table 8). Furthermore, a second olaparib-resistant cell line, SK-N-AS (non-*MYCNA*, TP53 wild-type)^19^ displayed relative resistance to combination olaparib + palbociclib treatments following DDR protein knock-down (Fig 5A, Supplementary Fig 6, Supplementary Table 8). Palbociclib single agent and combination treatments were associated with expected reductions in phosphorylated Rb (p-Rb) (Fig 5B). Interestingly, following siDDR protein knock-down, p-Rb levels were seen to increase compared to control siRNA treatment, consistent with observed activation of E2F-dependent signaling in *EGFP;MYCN;tp53^-/-^;*DDR-deficient zebrafish tumors (Fig 4, 5D,E). This effect was found to further increase following olaparib single agent treatment (Fig 5B), suggesting opposing effects of olaparib and palbociclib on targeted Rb inhibition in NB cells, which could explain the observed lack of consistent drug synergy *in vitro*.

**Figure 5.**
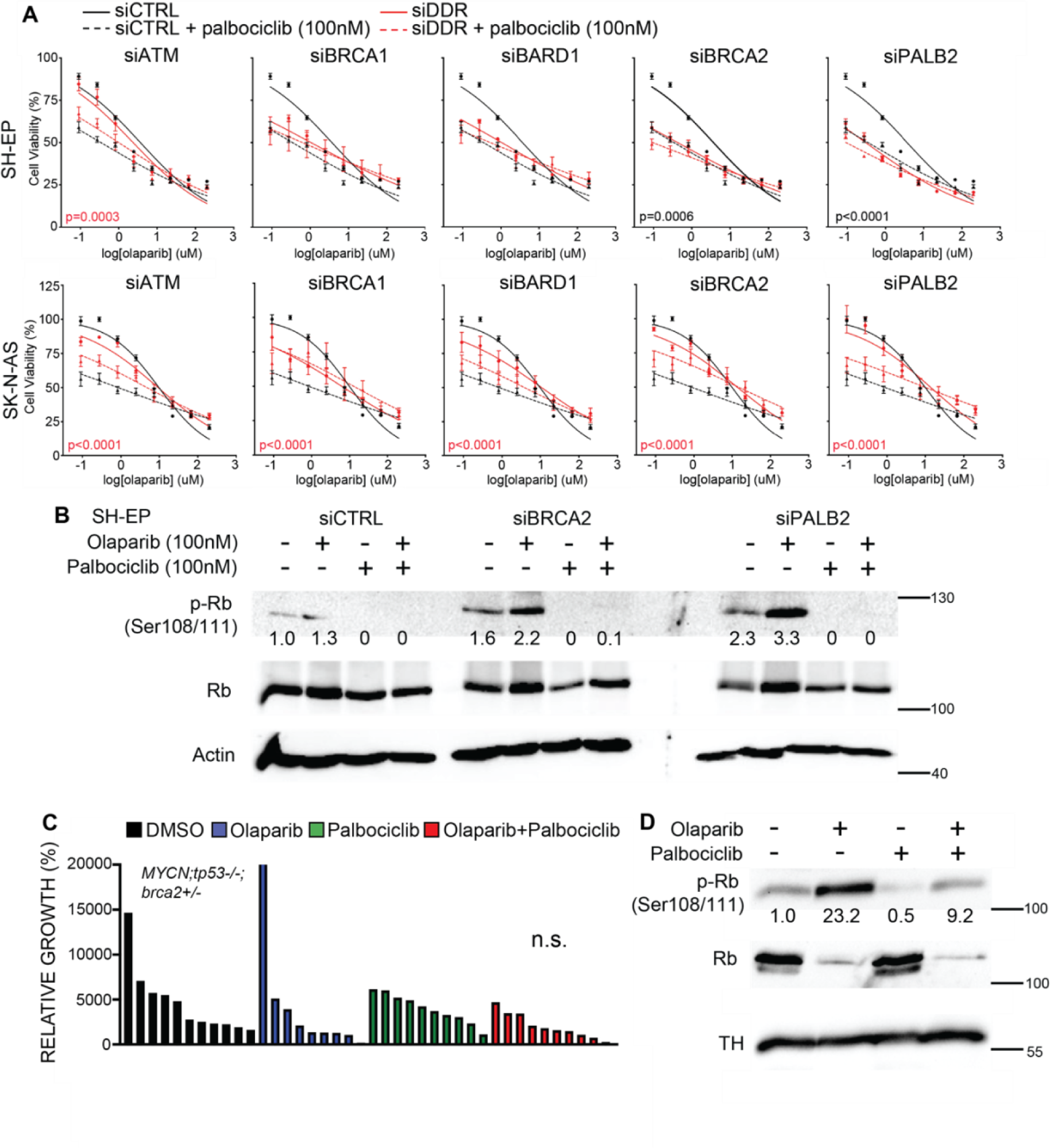
The CDK4/6 inhibitor palbociclib does not generally synergize with olaparib in DDR-deficient NB cells. **(A)** Cell viability (AlamarBlue) analysis in SH-EP and SK-N-AS human NB cell lines following siRNA-mediated gene knock-down of *ATM, BRCA1, BARD1, BRCA2,* and *PALB2*, and 72-hour treatment with olaparib [0.05-800uM] + palbociclib [100nM]. p-values represent comparisons of non-linear fits between combination olaparib + palbociclib treated siDDR and siControl (siCTRL) cells. Black p-values highlight significant increases in sensitivity (decreased IC_50_) with siDDR. Red p-values highlight significant resistance (increased IC_50_) with siDDR. **(B)** Western blot analysis of phosphorylated Rb (p-Rb Ser108/111) and total Rb protein in SH-EP cells in response to olaparib and/or palbociclib treatments following control (siCTRL), *BRCA2* (siBRCA2), and *PALB2* (siPALB2) knock-down. Actin is used as loading control. p-Rb is quantified below each lane and displayed as normalized first to total Rb protein followed by untreated control knock-down cells (siCTRL). **(C)** Quantification of relative growth after four weeks of dosing (4 consecutive days per week) DMSO vehicle control, olaparib (50mg/kg), palbociclib (100mg/kg) or olaparib + palbociclib to zebrafish engrafted with *EGFP;MYCN;tp53^-/-^;brca2^+/-^* NB (n=2 independent primary tumors). Differences between control and combination treatment groups is not significant (n.s.), Student’s t-test. **(D)** Western blot analysis of phosphorylated Rb (p-Rb Ser108/111) and total Rb protein in response to *in vivo* olaparib and/or palbociclib treatments. Tyrosine hydroxylase (TH) is used as a loading control for tumor cells following dissection at experimental end-point of bulk tumor. p-Rb is quantified below each lane and displayed normalized to total Rb protein and vehicle control-treated sample.

In parallel, we tested combination olaparib + palbociclib treatment on DDR-deficient NB *in vivo* following engraftment of *brca2* heterozygous mutant (*EGFP;MYCN;tp53^-/-^;brca2^+/-^*) zebrafish tumor cells into *prkdc^-/-^* immune-deficient hosts. In contrast to tumors with homozygous *brca2* loss (*EGFP;MYCN;tp53^-/-^;brca2^-/-^*) (Fig 3), tumors with heterozygous *brca2* gene knockout (*EGFP;MYCN;tp53^-/-^;brca2^+/-^*) were relatively resistant to olaparib treatments (50mg/kg olaparib, oral gavage 4 consecutive days/week) (Fig 5C). *EGFP;MYCN;tp53^-/-^;brca2^+/-^* NB growth was also not significantly inhibited by palbociclib treatment (100mg/kg palbociclib, oral gavage 4 consecutive days/week) (Fig 5C). Notably, olaparib + palbociclib combination treatment (50mg/kg olaparib + 100mg/kg palbociclib, 4 consecutive days/week) failed to inhibit NB growth, suggesting a lack of drug synergy in *EGFP;MYCN;tp53^-/-^;brca2^+/-^ in vivo*. In dissected tumor samples, we detected reduced p-Rb levels following palbociclib single-agent treatment, compared to vehicle control-treated tumors, while tumors from animals treated with olaparib displayed significant increases in p-Rb (Fig 5D). p-Rb was partially suppressed compared to olaparib treatment by combination treatment with palbociclib (Fig 5D), again suggesting opposing effects of olaparib and palbociclib on Rb when dosed together in DDR-deficient NB. While our results are consistent with palbociclib-mediated inhibition of DDR proficient NB growth *in vitro*^84,90,92^ (Fig 5A), we found that overall, palbociclib does not readily synergize with olaparib to inhibit DDR-deficient NB. Altogether, our results suggest that NB patients with DDR pathway alterations may be more resistant to palbociclib CDK4/6i therapy, especially in combination with PARPi, and that targeting alternative DNA damage repair and/or cell checkpoint pathways may be more optimal strategies.

### Targeting the G2M checkpoint pathway in combination with PARP inhibition in DDR-deficient neuroblastoma

Based on our transcriptomics data supporting ATR pathway activation in *brca2*^+/-^ and *atm^+/-^* zebrafish NB (Figure 4C, Supplementary Table 6), we next asked whether targeting the replicative stress response and G2M checkpoint using the ATR inhibitor (ATRi) ceralasertib (AstraZeneca; AZD6738) would be an effective strategy. This approach is further supported by the demonstrated efficacy of ATRi alone and in combination with PARPi in a number of adult cancers with mutations in DDR genes^93–96^. We first tested ceralasertib *in vitro* in combination with olaparib using SH-EP and SK-N-AS cells with siRNA-mediated knock-down of *ATM, BRCA1, BARD1, BRCA2,* and *PALB2* (Fig 6A). Remarkably, across both cell lines and target gene knock-downs (for all except for siATM SH-EP cells), ceralasertib led to increased sensitivities to olaparib, as shown by reductions in IC_50_ values, compared to control siRNA treated cells (Fig 6A, Supplementary Table 8). Ceralasertib treatment was associated with reductions in phosphorylated Chek1 (pChk1), a direct target and indicator of activated ATR (Fig 6B). Furthermore, in SK-N-AS cells with DDR protein knock-down, ceralasertib treatment led to increased DNA damage marked by y-H2AX, compared to control siRNA, and this effect was further enhanced by combination treatment with olaparib (Fig 6B). Importantly, combination drug treatments in SH-EP and SK-N-AS cells with DDR protein knock-down dramatically increased cleaved-PARP protein, indicating increased apoptosis and cell death, supporting a synergistic inhibitory effect of ceralasertib + olaparib on DDR-deficient human NB *in vitro*.

**Figure 6.**
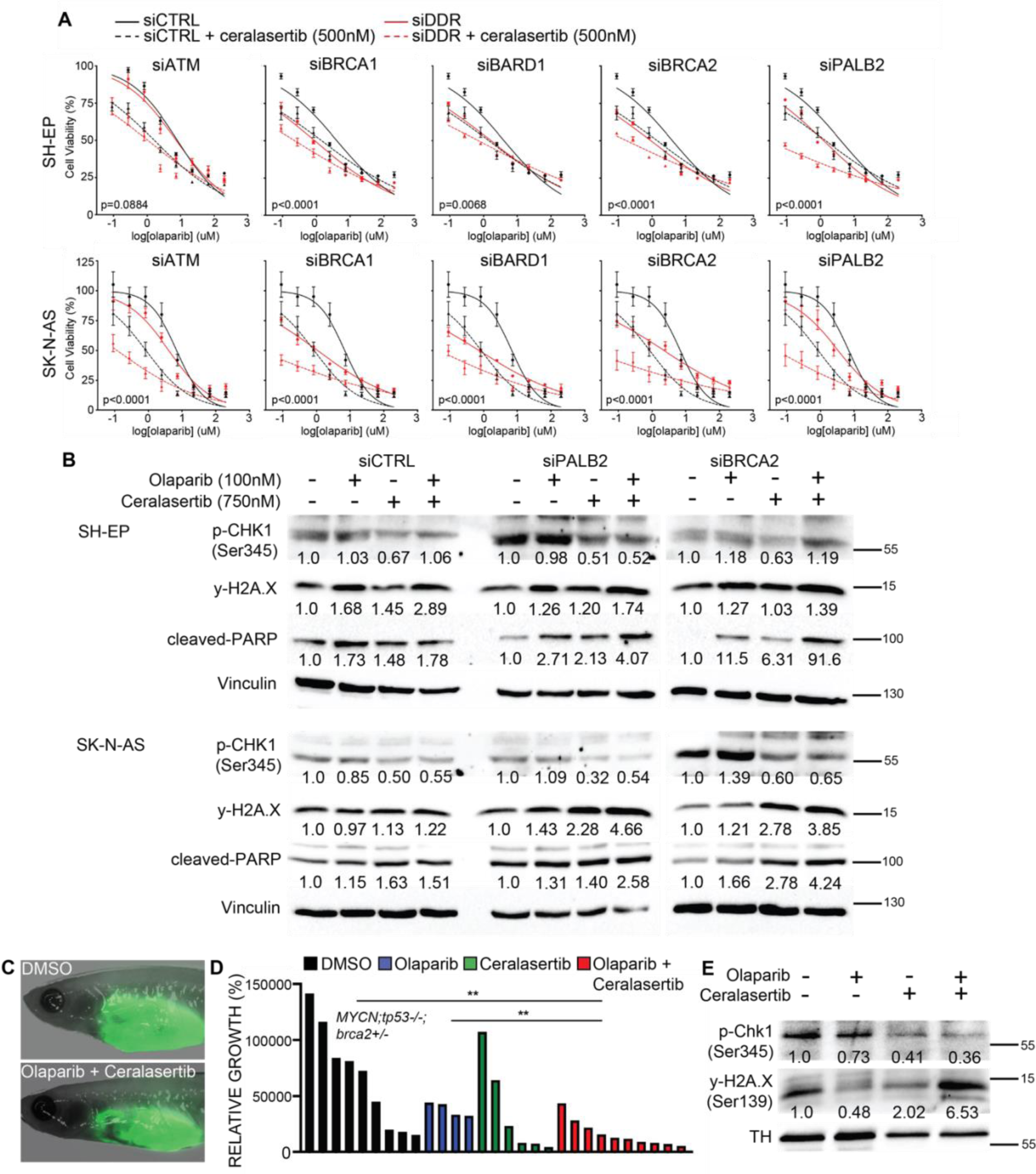
The ATR inhibitor ceralasertib enhances olaparib-mediated inhibition of DDR-deficient NB cells. **(A)** Cell viability (AlamarBlue) analysis in SH-EP and SK-N-AS human NB cell lines following siRNA-mediated gene knock-down of *ATM, BRCA1, BARD1, BRCA2,* and *PALB2*, and treatment with olaparib [0.05-800uM] + ceralasertib [500nM]. p-values represent comparisons of non-linear fits between olaparib + ceralasertib treated siDDR and siControl (siCTRL) cells. **(B)** Western blot analysis of phosphorylated CHK1 (pCHK1, Ser345), phosphorylated H2A.X (Ser345, y-H2A.X), and cleaved-PARP in SH-EP and SK-N-AS cells following control (siCTRL), *BRCA2* (siBRCA2), and *PALB2* (siPALB2) knock-down. Vinculin is used as loading control. Relative protein levels are quantified below each lane and displayed normalized to Vinculin followed by vehicle control drug-treated cells within each siRNA treatment. **(C)** Live *casper;prkdc^-/-^* immune-deficient zebrafish engrafted with *EGFP;MYCN;tp53^-/-^;brca2^+/-^*at experimental endpoint following four weeks of dosing DMSO vehicle control and combination olaparib (50mg/kg, 4 consecutive days/week) + ceralasertib (40mg/kg, 4 consecutive days/week) treatments. **(D)** Quantification of relative growth after four weeks of dosing (4 consecutive days per week) DMSO vehicle control, olaparib (50mg/kg), palbociclib (40mg/kg), and combination olaparib + ceralasertib to zebrafish engrafted with *EGFP;MYCN;tp53^-/-^;brca2^+/-^* NB. Combined results from 2 independent primary tumors, minimum 2 technical replicates for each treatment cohort. Differences between control and combination treatment group is significant (p=0.0037), Student’s t-test. **(E)** Western blot analysis of phosphorylated Chk1 (p-Chk1, Ser345), and y-H2A.X (Ser139) in response to *in vivo* olaparib and/or ceralasertib treatments. Tyrosine hydroxylase (TH) is used as a loading control for tumor cells following dissection at experimental end-point. Relative p-Chk1 and y-H2A.X levels are quantified below each lane and displayed normalized to TH protein and vehicle control-treated sample.

We also tested combination olaparib + ceralasertib treatment on *brca2* heterozygous (*EGFP;MYCN;tp53^-/-^;brca2^+/-^*) zebrafish NB cells engrafted into *prkdc^-/-^* immune-deficient hosts. As single agents, olaparib (50mg/kg olaparib, 4 consecutive days/week) and ceralasertib (40mg/kg, 4 consecutive days/week) failed to significantly inhibit *EGFP;MYCN;tp53^-/-^;brca2^+/-^* growth *in vivo*. However, combination olaparib + ceralasertib treatment (50mg/kg olaparib + 40mg/kg ceralasertib, 4 consecutive days/week) led to a significant reduction of tumor growth, compared to vehicle-control after 4 weeks of treatments (Fig 6C,D). Furthermore, combination treatment led to reductions in pChk1 protein in bulk tumor lysates, as well as increased y-H2AX (Fig 6E), suggesting on target drug effects and increased DNA damage, incompatible with optimal tumor cell viability *in vivo*. Altogether, our results suggest an enhanced molecular vulnerability of DDR-deficient NB to combination olaparib PARPi + ceralesertib ATRi, and a potential therapeutic strategy for patients with alterations in DDR genes.

## Discussion

This is the first *in vivo* modeling study of DDR loss-of-function in NB. We reveal functional roles for DDR proteins on NB tumorigenesis, metastasis, gene expression signatures, and drug sensitivities *in vivo* as well as in human cells. Using high-throughput mutagenesis approaches, we were able to assess the impact of multiple DDR genes, including *Brca2, Palb2, Atm,* and *Bard1*, and gain a better understanding of a role for DDR-deficiency in enhancing NB formation as well as metastasis in patient with tumors harboring alterations in DDR genes. We also demonstrate that the gene expression profiles of these tumors may provide important rationale for PARPi therapy and PARPi + ATRi for the treatment of molecularly selected patients with NB and homozygous or hemizygous DDR pathway alterations, respectively. Collectively, our study highlights both the feasibility and utility of modeling patient-relevant loss-of-function DDR pathway mutations, for the purposes of mechanistic discovery as well as validating certain genetic determinants as biomarkers for targeted therapies. These findings may also impact insights into genetic predisposition variants and surveillance approaches. Pathogenic and likely pathogenic DDR pathways variants have also been identified in children and adolescents with malignancies such as CNS tumors and sarcomas. Thus, our findings support further functional and mechanistic studies to define potentially more generalizable mechanisms of DDR-dependent tumorigenesis in different pediatric cancers.

Previous implications for roles for DDR pathways in NB have been made from sequencing tumors as well as germline DNA from pediatric patients^20–23^. Furthermore, recent descriptions of mutational signature SBS3 in pediatric solid tumors, including NB, provide evidence for a role for DDR variants in driving oncogenic processes and tumor evolution^20^. Here we show that loss-of-function mutations in *brca2*, *atm,* and *palb2* enhance *MYCN;tp53^-/-^* NB, leading to aggressive phenotypes in zebrafish, providing *in vivo* evidence of a functional role for DDR-deficiency in NB initiation and progression. Independent findings in isogenic NB cells with and without expression of the *BARD1* (BRCA1 associated ring domain protein 1) gene provide additional support for a functional role for BARD1 in mediating DNA repair, highlighting the impact of loss-of-function *BARD1* variants^37^. Future investigations will be important to further assess DNA repair signatures associated with different DDR mutations and potentially identify conserved loci affected, to better understand underlying mutational mechanisms driving high-risk phenotypes including metastasis.

*BRCA1, BRCA2, PALB2* and *BARD1* have been broadly studied in adult cancers and targeting DDR-deficiency with PARPi is an effective therapeutic strategy for certain adult patients with pathogenic germline or somatic variants in these genes^28,31,68^. For germline DDR mutations in adults, somatic loss-of-heterozygosity (LOH) is often seen in tumor cells, with data suggesting roles for LOH in *BRCA-*associated cancer subtypes including breast, ovarian, pancreatic and prostate cancers^26,27,97^. However, while LOH of *PALB2* was detected by sequencing in one of our NB tumors, LOH is not commonly found among pediatric cancer patients with germline HR variants^20,98^. Furthermore, results from adult cancer patients with prostate cancer indicate PARPi sensitivities in the absence of LOH^99^, altogether suggesting that a second genetic “hit” at the same single DDR gene locus is not necessarily required for tumorigenesis and/or targeted drug responses. In many pediatric solid tumor samples multiple somatic mutations affecting different DDR genes were detected^20^, suggesting that collaborating loss-of-function mutations may affect the pathogenesis of certain NBs and other embryonal tumors. Inherited haploinsufficiency of certain DDR loss-of-function alleles was sufficient to increase tumor formation, metastasis, and molecular signatures indicative of DNA repair defects and aggressive tumor cell behaviors in our zebrafish models. LOH at the *brca2* locus was not detected in 3/3 germline heterozygous tumors analyzed (data not shown), and together with patient genetic data as well as recent functional investigations into *BARD1* haploinsufficency in human NB cells resulting in DNA repair defects^20,37^, our functional and sequencing data suggests that LOH as not necessarily required at a single DDR gene locus for the NB-associated effects observed. Moreover, in a recent study that identified germline pathogenic variants in *BARD1* enriched in patients with neuroblastoma, there was no evidence of LOH or bi-allelic inactivation^22^. Interestingly, these *BARD1* variants were more common in patients with high-risk metastatic disease, which supports our findings of enhanced metastases in DDR-deficient zebrafish. Future investigations will be needed to assess the presence and/or role for additional pathway “hits” as well as potential non-coding variants *in vivo*.

ATM is a major DDR protein involved in sensing double strand DNA breaks and 30-40% of NBs display losses of chromosome 11q, which includes *ATM* (11q22.3)^100,101^. *ATM* loss is associated with inferior survival independently of *MYCNA* status and has previously been shown to affect NB cell growth and PARPi sensitivity in human NB cells^33–35,102,103^. Here, we functionally validate that *atm* loss leads to a highly penetrant and aggressive NB phenotype in zebrafish, and *atm* haploinsufficiency (*atm^+/-^*) is sufficient to enhance penetrance and metastasis of *MYCN;tp53^-/-^* zebrafish NB, supporting a major role for *ATM* loss in 11q deletions and previous evidence highlighting therapeutic potentials in ATM-deficient NB^34,35,104^. Although 11q loss and *MYCNA* are genetically distinct NB cohorts and rarely co-occur, mechanistic associations including MYCN/ATM feedback loops suggest that loss or epigenetic downregulation of ATM may collaborate with MYCN during cellular transformation^103,105^. Further investigations into signaling downstream of MYCN, ATM, and/or other DDR signaling components will help clarify specific collaboration/cooperation between MYCN activity and ATM loss, specific *in vivo* effects of ATM hemizygous compared to homozygous loss-of-function mutations in NB^106^, and conserved molecular effects involved in high-risk phenotypes including metastasis, across NB genetic cohorts.

PARP inhibitors (olaparib, talazoparib) and ATR inhibitors (ceralasertib, elimusertib) have shown efficacy as single agents, but increasingly with other agents including immune checkpoint inhibitors^107,108^. For adult patients with breast, ovarian and prostate cancers, indications for these drugs and clinical benefit have been shown to be dependent on germline and/or somatic alterations of certain DDR genes, including but not limited to *BRCA1/2*, *PALB2*, and *ATM*^30–32^. More recently, the spectrum of tumors with documented responses to PARP inhibition has expanded, although to date, most published pediatric trials of PARP inhibitors or other drugs have not required DDR mutations as a selection or eligibility criteria. Precision pediatric precision basket trials such as the NCI MATCH and ITTC e-SMART are prospectively assigning patients to PARP inhibitor therapies alone or in combination with other agents for patients whose tumors have alterations in certain subsets of DDR variants^109–111^. Combination PARPi + ATRi was previously shown to inhibit NB growth due to increased replicative stress responses in NB cells^112^. Our transcriptomic data suggests even further enhancement of replicative-stress associated pathway activation in NB tumors with DDR pathway mutations, supporting DDR variants as eligibility for G2M checkpoint and/or DDR pathway inhibitor-based regimens for NB. Furthermore, our models provide opportunities to functionally characterize additional and less common variants identified through retrospective and prospective next generation sequencing studies. Genomic studies of pediatric tumors have identified variants in genes implicated in DDR, with many yet understudied in the context of pediatric cancer such as *CHEK2*, *RAD51* and multiple Fanconi complex proteins^20–25^. Examining roles for patient-derived loss-of-function variants in these genes in zebrafish NB models could provide additional evidence for their pathogenicity and the identification of pre-clinical targets and thus, for their inclusion in precision trials.

In addition to therapeutic relevance, our findings have important impacts on our understanding of the role of DDR variants in pediatric cancer predisposition syndromes. Currently, most pediatric genetic predisposition syndrome programs consider “adult-onset cancer” variants as not impacting needs for tumor surveillance during childhood. Increasingly, germline and somatic DDR variants similar to those identified in our KiCS program are detected in patients with pediatric tumors including NB, sarcomas and CNS tumors^20,22,113–117^. However, based on epidemiologic data and lack of mechanistic studies it has not been clear whether these variants play a causal role or are coincidental passenger variants in pediatric cancer pathogenesis. A recent meta-analysis of eleven publications of germline testing concluded that heterozygous mutations in *BRCA1, BRCA2, PALB2, CHEK2, MSH2, MSH6, MLH1,* and *PMS2* in pediatric patients do contribute to cancer risk in this population but with reduced penetrance; therefore, predictive genetic testing and/or enhanced surveillance is not currently recommended^118^. However, notably the specific impact on tumor subtypes may not be apparent in these pooled analyses, especially given that many of these variants are rare. In contrast, a large recent study focused on germline sequencing in NB patients reported increased risk linked to germline variants in *BARD1*^22^. Data from our models support roles for additional pathogenic/likely pathogenic “adult-onset cancer” DDR variants in NB tumorigenesis and metastasis, highlighting the need for consideration of DDR gene variants in future pediatric surveillance and clinical pediatric oncology practice.

Taken together our findings provide important evidence for a role for DDR gene variants in NB formation and metastasis *in vivo*. Furthermore, sequencing studies and gene expression analyses provide mechanistic insights into downstream signaling pathways, including the potential to exploit specific cellular vulnerabilities including ATR-mediated checkpoint activation in combination with PARP inhibitors, to treat patients with NB with alterations in certain DDR pathway genes.

## Materials and Methods

### Zebrafish lines and maintenance

Zebrafish were raised and maintained according to standard procedure. All studies, maintenance and monitoring were performed in accord with animal use protocols #1000054111 and #1000064586, approved by the Hospital for Sick Children Animal Care Committee. Zebrafish lines *Tg(dbh:EGFP)* and *Tg(dbh:EGFP-MYCN)* were previously described and a kind gift from Shizhen Zhu (Zhu et al., 2012). They are designated *EGFP* and *MYCN* in the text, respectively. CG1*tp53^del^* were previously described (Ignatius et al., 2018), and designated *tp53^-/-^* in the text. *casper;prkdc^D^*^3612^*^fs^* (*prkdc^-/-^*, SCID) zebrafish were previously described (Moore et al., 2016), and a kind gift from David Langenau. *Brca2^hsc^*^195^, *atm^hsc^*^196^ and *palb2^hsc^*^197^ mutant alleles were generated using CRISPR/Cas9 (see below) and designated *brca2^-/-^*, *atm^-/-^* and *palb2^-/-^* when homozygous in the text.

### CRISPR/Cas9 Gene Targeting

CRISPR/Cas9 target sites were selected for *brca2, atm, palb2* and *bard1* against the danioRer11/GRCz11 genome assembly using CHOPCHOP v3 (Labun et al., 2019). Individual target sites are listed in Table S2. *In vitro* transcription of each gRNA was performed using the EnGen sgRNA Synthesis Kit (NEB #E3322), according to manufacturers directions. gRNAs were purified using the Monarch RNA Cleanup Kit (NEB), according to manufacturers recommendations. 50pg of each gRNA (Table S1) was co-injected with 1ng purified Cas9 protein (PNA Bio, CP01) into *EGFP;MYCN;tp53^-/-^* embryos at the one-cell stage. Targeting efficiency of each gRNA was assessed prior to co-injection with other gRNAs targeting the same gene using DNA extraction at 1-3 dpf, followed by PCR-based amplification of target sequence using genotyping primers listed in Table S2. PCR fragments were submitted for Sanger sequencing (Centre for Applied Genomics, Hospital for Sick Children). Raw .ab1 sequencing files were assessed and compared to un-injected control using ICE Analysis (Synthego Performance Analysis, v3.0, ice.synthego.com/#/) and Indigo analysis (GEAR-Genomics, gear-genomics.com, ^119^). Germline transmission was assessed by genotyping the progeny of mosaic-injected animals outcrossed to *tp53^-/^*^-^. The genotypes of all subsequent generations were assessed using DNA extracted from individual fin clips serving as template using the primers listed in Table S2. Products were validated using Sanger sequencing at the Centre for Applied Genomics, Hospital for Sick Children, or visualized using standard gel electrophoresis for routine genotyping.

### Mosaic CRISPR/Cas9 Injected NB Cell Analysis

Viable EGFP+ NB cells from one of each *brca2, atm, palb2* and *bard1* mosaic CRISPR/Cas9-injected *EGFP;MYCN;tp53^-/-^* zebrafish were sorted (DAPI-negative/EGFP+) at the SickKids-UHN Flow Cytometry Core Facility on a Sony MA900 VBYR cell sorter. Cells were lysed for DNA and PCR-based genotyping was performed surrounding target sites, followed by Sanger sequencing, ICE analysis (Synthego Performance Analysis, v3.0, ice.synthego.com/#/), and/or Indigo analysis (gear-genomics.com, ^119^). For *brca2* and *palb2* 5’ and 3’ primer combinations (Table S2) were used to capture all possible deletion fragments.

### Neuroblastoma tumor detection

Mosaic primary injected and/or stable zebrafish lines were observed for evidence of EGFP+ fluorescent tumors starting at 6 wpf. Fish with tumors were euthanized after anesthesia and characterized by histopathological hematoxylin and eosin (H&E) staining and immunohistological analysis or immunofluoresence using antibodies against PCNA (Cell Signaling, D3H8P), tyrosine hydroxylase (Pel-Freez Biologicals, P40101-150), EGFP (Abcam, ab6673), and y-H2A.X (Abcam, ab228655). IHC was performed using DAB procedure (Signal Stain Boost IHC Detection Reagent, CS#8114), according to manufacturers recommendations. EGFP immunofluorescence staining was performed using Invitrogen Alexa Fluor 555 donkey anti-goat IgG (A21432).

### DNA Extraction, Whole Genome Sequencing and Variant Analysis

Tumor burdened zebrafish were sacrificed at 4-5 months of age, and tumors macrodissected. EGFP+ tumor cells and EGFP-negative non-tumor cells were dissociated and sorted at the SickKids-UHN Flow Cytometry Core Facility on a Sony MA900 VBYR cell sorter. Cells were pelleted at 1000xg and lysed for DNA using the Qiagen DNeasy Blood and Tissue Kit, as per manufacturer instructions. Total genomic DNA was PCR amplified and processed for whole genome sequencing using Illumina NovaSeq at the Hospital for Sick Children’s Centre for Applied Genomics (TCAG). 150bp paired-end reads at a sequencing depth of 60x was used for tumor DNA and 30x for non-tumor tissue.

FASTQ files were aligned to the GRCz11/danRer11 reference genome using BWA-mem (v0.7.8). Duplicates were marked with Sambamba (v0.7.0). Small substitution and indel calls were made using MuTect2 (v2.2) from GATK (v4.1.9). Somatic substitutions (SNVs and indels) were excluded using the following rules: failed any of internal MuTect2 filters, tumor depth at the position <20, variant allele depth in tumor <5, variant allele depth in germline ≥1, and VAF in tumor <0.1. VEP (Variant Allele Predictor; v111) with dbSNP (v150), refseq (v106.20170823 - GCF_000002035.6) and sift (v6.2.1) were used to annotate the SNV and indel variants^120^. Structural variants (deletions, duplications, inversions, and translocations) were called using manta (v1.6.0)^121^. SVs that failed any of the Manta’s internal filters or had IMPRECISE breakpoints were excluded.

### Bulk RNAseq Library Preparation, Quantification and Differential Gene Expression Analysis

Tumor burdened zebrafish were sacrificed, and tumors macrodissected. EGFP+ tumor cells were dissociated and sorted from bulk tissue at the SickKids-UHN Flow Cytometry Core Facility on a Sony MA900 VBYR cell sorter, before pelleting and lysis using Qiagen RNeasy Mini Kit, as per manufacturer instructions. Sequence ready polA-enriched libraries were prepared using the NEB Ultra II Directional mRNA prep kit for Illumina (NEB, E7760). Paired-end sequencing at a a targeted depth of ∼50,000 million reads/sample was performed using the Illumina NovaSeq platform at the Centre for Applied Genomics (TCAG). Raw .fastq data was processed using Salmon quantification of transcripts. A “decoy-aware” index was built with the *Danio rerio* transcriptome and genome using the GRCz11 assembly with a k-mers length of 23. Samples were quantified with the following arguments: -r, --seqBias, --mp −3, --validateMappings, -- rangeFactorizationBins 4. Normalized counts outputs from DESeq2 were utilized for GSEA analysis without any further trimming or processing, as recommended by the GSEA user guide^122,123^. Zebrafish transcripts were assigned known or high-confidence Human orthologs using Ensembl BioMart. Expression signatures were compared against Hallmark and curated gene sets from the Molecular signatures database, as well as DNA repair signature from Knijnenburg et al.^124,125^

### Drug Dosing and Analysis

Primary NB was dissected from *EGFP;MYCN;tp53^-/-^;brca2^-/-^, EGFP;MYCN;tp53^-/-^;brca2^+/-^*, and *EGFP;MYCN;tp53^-/-^;brca2^+/+^* zebrafish at 18-30 weeks. Bulk tumor tissue was dissociated into a single cell suspension and injected into the interperitoneal space of 3-4 month old *casper;prkdc^-/-^* immune deficient zebrafish^71^. Epifluorescent whole-animal imaging was performed seven days post injection and during the drug course, using the same UV light intensity and camera exposure for each primary tumor (Zeiss Axiozoom V16). Tumor size was determined by quantifying 2D image area using Image J and multiplying by the average GFP fluorescence intensity, as previously described^72,126^. Engrafted fish were orally gavaged with 10uL of single agent or combination as indicated of olaparib (AbMole, AZD2281), TMZ (AbMole, M2129), palbociclib (Selleckchem, S1579), and/or ceralasertib (Selleckchem, S7693) using a Hamilton 22 G needle fitted with 22 G soft-tip catheter tubing. Drugs were administered weekly for olaparib + TMZ, or for 4 consecutive days per week for up to four weeks or to humane endpoint. Animals were sacrificed and fixed in 4% paraformaldehyde, sectioned, and examined histologically using PCNA (Cell Signaling, D3H8P) and Cleaved-caspase 3 staining (Cell Signaling, Asp175, 5A1E). Tumors were dissected and lysed for protein detection by Western blot using antibodies against phospho-Rb Ser807/811 (Cell Signaling, #9308), Rb (GeneTex, GTX54191), phospho-Chk1 Ser345 (Cell Signaling, 133D3, #2348), and tyrosine hydroxylase (Pel-Freez Biologicals, P40101-150).

### *In vitro* olaparib, palbociclib and ceralasertib Treatments

Neuroblastoma cell lines SH-EP, IMR-32 and SK-N-AS were purchased from ATCC, tested for *Mycoplasma* (InvivoGen) and authenticated using short tandem repeat (STR) analyses (TCAG Sequencing Facility, Toronto, ON). IMR-32 cells were cultured in EMEM (Wisent) supplemented with 10% FBS (Wisent), 0.1mM nonessential amino acids (Gibco), and 1mM sodium pyruvate (Gibco). SH-EP and SK-N-AS cells were cultured in RPMI medium (Wisent) supplemented with 10% FBS, 3mM L-glutamine (Gibco), and 1mM sodium pyruvate (Gibco). For DDR-knockdown assays, SH-EP, IMR-32, and SK-N-AS cells were transfected with 150pM of siRNA (Silencer Select siRNAs, ThermoFisher Scientific) using RNAiMAX lipofectamine reagent (Invitrogen). 48 hours post-transfection, 3-5 x 10^3^ cells were seeded in 96-well plates and treated with olaparib (MedChemExpress, #HY-10162), palbociclib (Selleckchem, S1579), and ceralasertib (Selleckchem, S7693) for 72-96 hours. AlamarBlue reagent (Invitrogen) was added for 16-18 hours, and fluorescence was assessed using a microplate reader (Spectra MAX Gemini EM, Molecular Devices) with a l540 excitation/l590 emission filter. IC_50_ curves were calculated using GraphPad Prism 9 software (GraphPad Software Inc). Experiments included 3 technical replicates and were independently performed twice. Cell lysates were collected and assessed using Western blot and antibodies against ATM (Cell Signaling, D2E2, #2873), BARD1 (Santa Cruz, E-11, sc-74559), BRCA1 (Cell Signaling, ABX9F, #14823), BRCA2 (Cell Signaling, D9S6V, #10741), PALB2 (Cell Signaling, E9R2W, #30253), phospho-Rb Ser807/811 (Cell Signaling, #9308), Rb (Cell Signaling, 4H1, #9309) GTX54191), phospho-Chk1 Ser345 (Cell Signaling, 133D3, #2348), cleaved-PARP (Cell Signaling, D214, #9541), y-H2A.X (ab228655), Vinculin (Millipore, V284, 05-386) and/or Actin (Cell Signaling, #4968).

## Acknowledgements

We thank Dr. Shizhen Zu for Tg(*dbh:*EGFP-MYCN) and Tg(*dbh:*EGFP) zebrafish strains, and Dr. David Langenau for CG1*tp53^del^* (*tp53^-/-^*) and *casper;prkdc^-/-^* zebrafish. We also thank the Centre for Applied Genomics (TCAG) for sequencing, the SickKids-UHN Flow Cytometry Facility for cell sorting, the Centre for Phenogenomics (TCP) Histopathology core for tissue processing and staining, and the Zebrafish Genetics and Disease Models Facility management and support team for zebrafish husbandry.

---

**Supplementary Figure 1.**
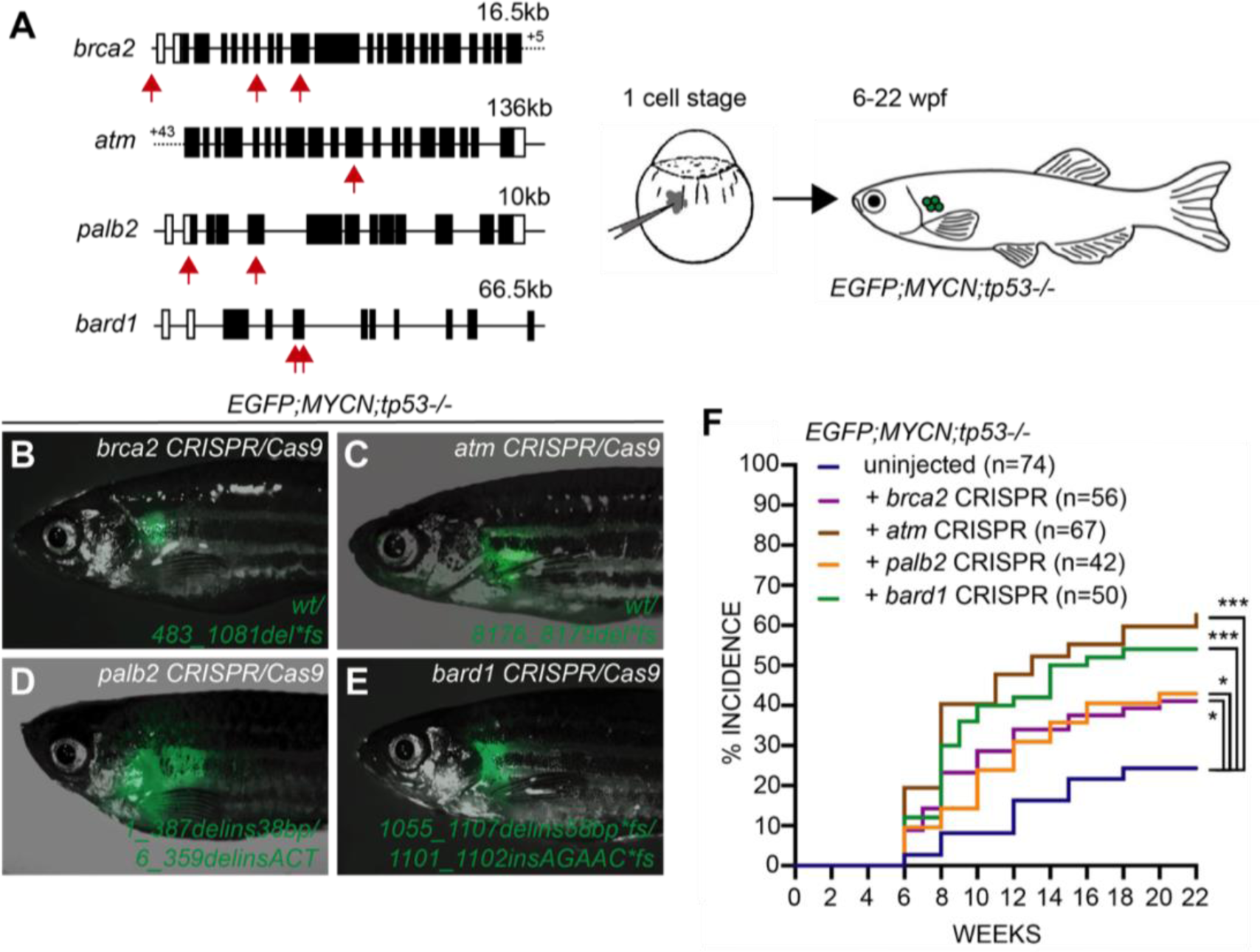
Mosaic (somatic) targeting of DDR pathway genes increases the penetrance of MYCN-induced neuroblastoma. **(A)** gRNA target sites and injection strategy for CRISPR/Cas9-mediated mutagenesis of *brca2, atm, palb2* and *bard1*. Gene exons are represented by black boxes and solid lines represent introns. gRNAs were co-injected with Cas9 protein into *EGFP;MYCN;tp53^-/-^*embryos at the 1-cell-stage, followed by assessment of EGFP+ NB formation, starting at 6 weeks post fertilization (wpf) up to 22 weeks. **(B-E)** EGFP expression in the interrenal gland of 12-15 wpf *EGFP;MYCN;tp53^-/-^* zebrafish injected at the one-cell stage with sgRNAs/Cas9 targeting *brca2* (B), *atm* (C), *palb2* (D), and *bard1* (E). Gene targeting was verified in sorted EGFP+ NB cells, with detected gene mutations indicated in green. *wt*, wild-type at all target sites. **(F)** Kaplan-Meier curves of NB frequency in un-injected *EGFP;MYCN;tp53^-/-^* zebrafish and *EGFP;MYCN;tp53^-/-^* zebrafish injected with sgRNAs/Cas9 targeting *brca2*, *atm*, *palb2*, and *bard1*. Significant increases in EGFP+ NB incidence was detected for primary mosaic injected *brca2* (p=0.0235, n=56), *atm* (p<0.0001, n=67), *palb2* (p=0.0271, n=42), *bard1* (p=0.0002, n=50), compared to *EGFP;MYCN;tp53^-/-^*. *p<0.05, ***p<0.001, Log-rank test.

**Supplementary Figure 2.**
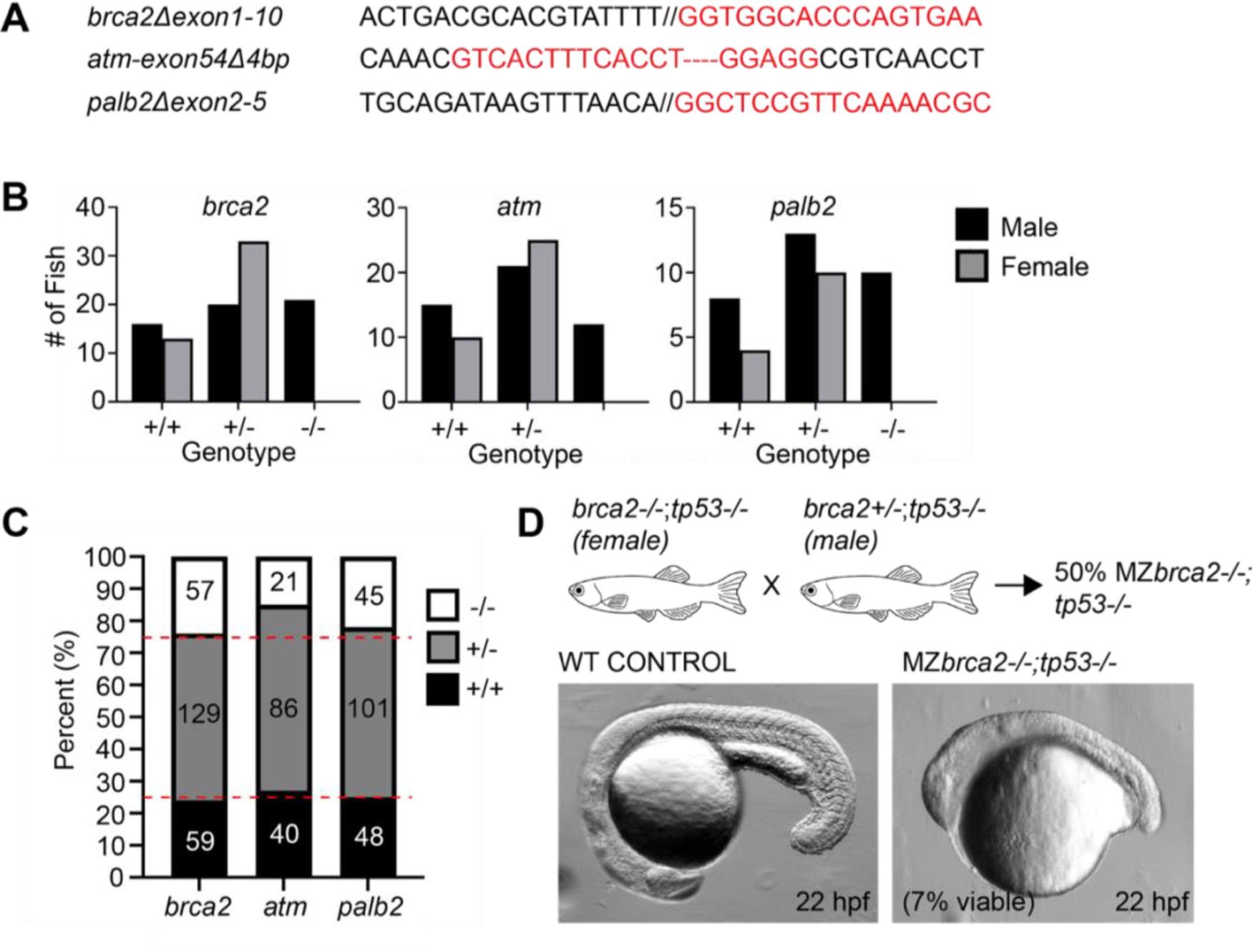
Isolation and characterization of *brca2, atm,* and *palb2* mutant lines. **(A)** Nucleotide sequences of *brca2Δexon1-10* (deletion between sequence 5’ upstream of exon 1 and exon 10), *atm-exon54Δ4bp* (4 base pair deletion in exon 54), and *palb2Δexon2-5* (deletion between exon 2 and 5) alleles surrounding CRISPR/Cas9 cleavage sites. Break points between distant exons are indicated by forward slashes and single nucleotide deletions are indicated by red dashes. **(B)** Quantification of male and female *brca2*, *atm,* and *palb2* wild-type (+/+), heterozygous (+/-), and homozygous (-/-) animals in *tp53^+/+^* and/or *tp53^+/-^* backgrounds. Homozygous DDR mutant zebrafish are strongly biased towards the male sex. **(C)** Proportion of *tp53^-/-^;brca2*, *tp53^-/-^;atm,* and *tp53^-/-^;palb2* wild-type (+/+), heterozygous (+/-), and homozygous (-/-) animals. All genotypes are viable up to 22 weeks of age, except for *tp53^-/-^;atm^-/-^* that display early lethality starting at 1 month post fertilization. Red dotted lines indicate expected Mendelian ratios. **(D)** Scheme of breeding strategy to achieve maternal and maternal zygotic (MZ) *brca2* mutant embryos. Homozygous *tp53^-/-^;brca2^-/-^* female zebrafish are only partially fertile (∼7-10% of embryos developed beyond the one-cell stage), with a small proportion of embryos viable up to 22 hpf, albeit with significant defects.

**Supplementary Figure 3.**
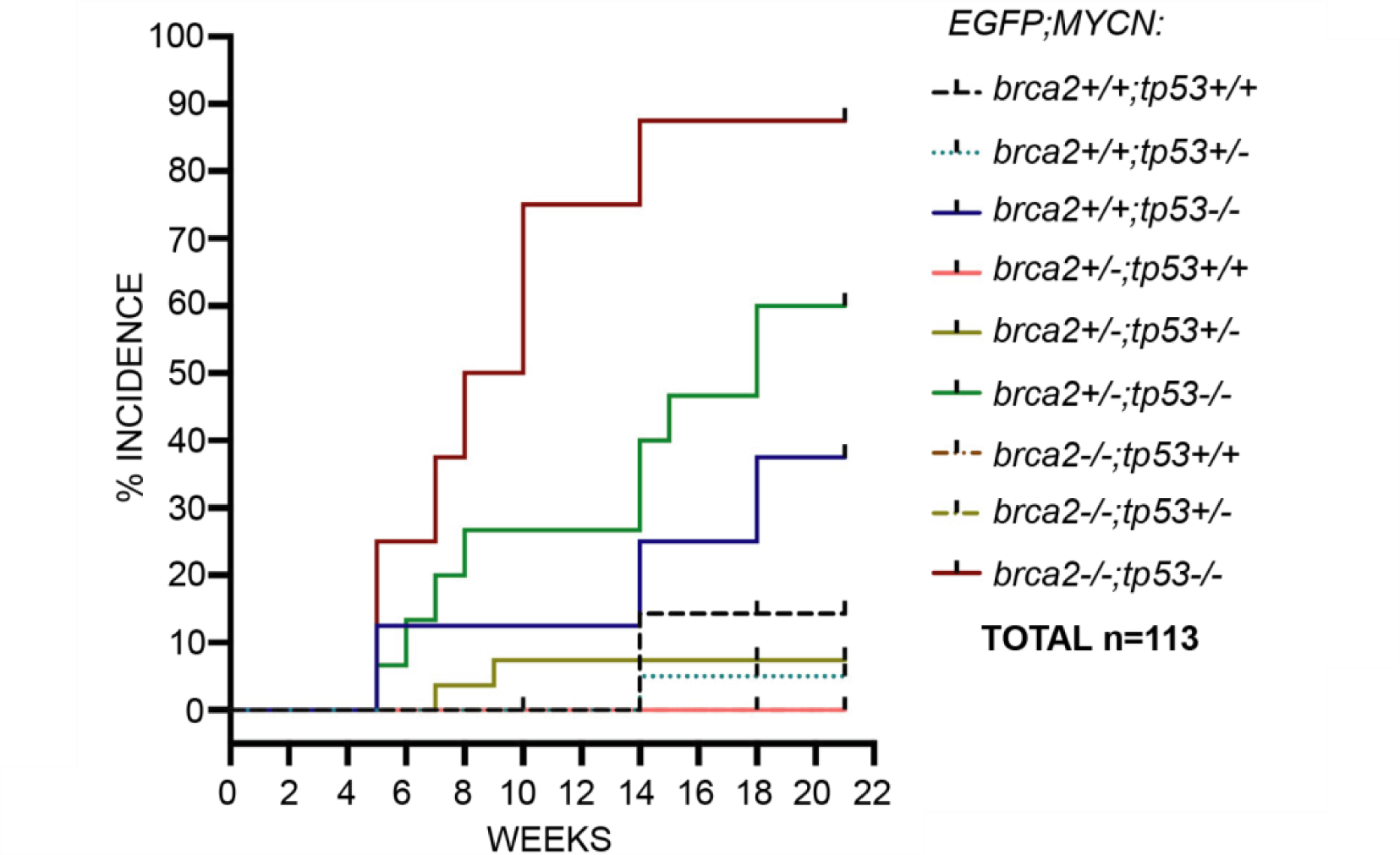
DDR-deficiency in loss-of-function *tp53* zebrafish enhances MYCN-induced NB formation. Cumulative frequency of NB onset among *EGFP;MYCN-* positive progeny from two independent *EGFP;MYCN;tp53^+/-^;brca2^+/-^* x *tp53^+/-^;brca2^+/-^* genetic crosses. NB formation is <20% at 22 weeks in *EGFP;MYCN* cohorts with wild-type or heterozygous *tp53* alleles.

**Supplementary Figure 4.**
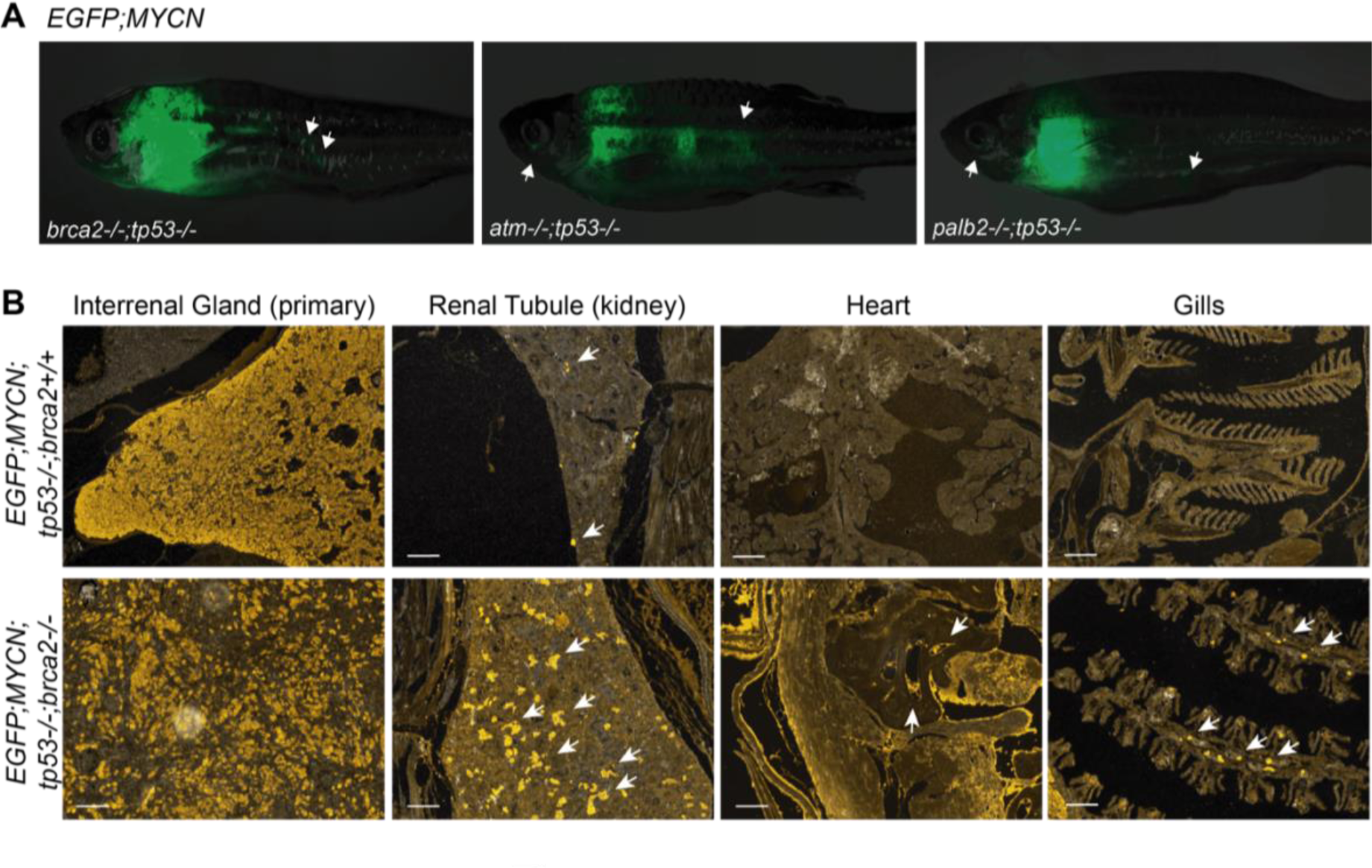
Metastasis in MYCN/DDR-deficient zebrafish models. **(A)** Disseminated EGFP+ NB in *EGFP;MYCN;tp53^-/-^;DDR* homozygous mutant zebrafish at 10-22 weeks. EGFP+ lesions visible on whole animals are indicated with white arrows. **(B)** Immunofluorescence staining in *EGFP;MYCN;tp53^-/-^* and *EGFP;MYCN;tp53^-/-^;brca2^-/-^* NB. Anti-EGFP immunofluorescence staining reveals EGFP+ NB nests of cells in the interrenal glands (primary initiation site) and renal tubules of tumor-burdened *EGFP;MYCN;tp53^-/-^*and *EGFP;MYCN;tp53^-/-^;brca2^-/-^* zebrafish. EGFP+ cells were also detected outside of the interrenal gland and renal tubules in *EGFP;MYCN;tp53^-/-^;brca2^-/-^*zebrafish prior to 22 wpf, including in the gills or within the heart chambers. Scale bar = 100μm.

**Supplementary Figure 5.**
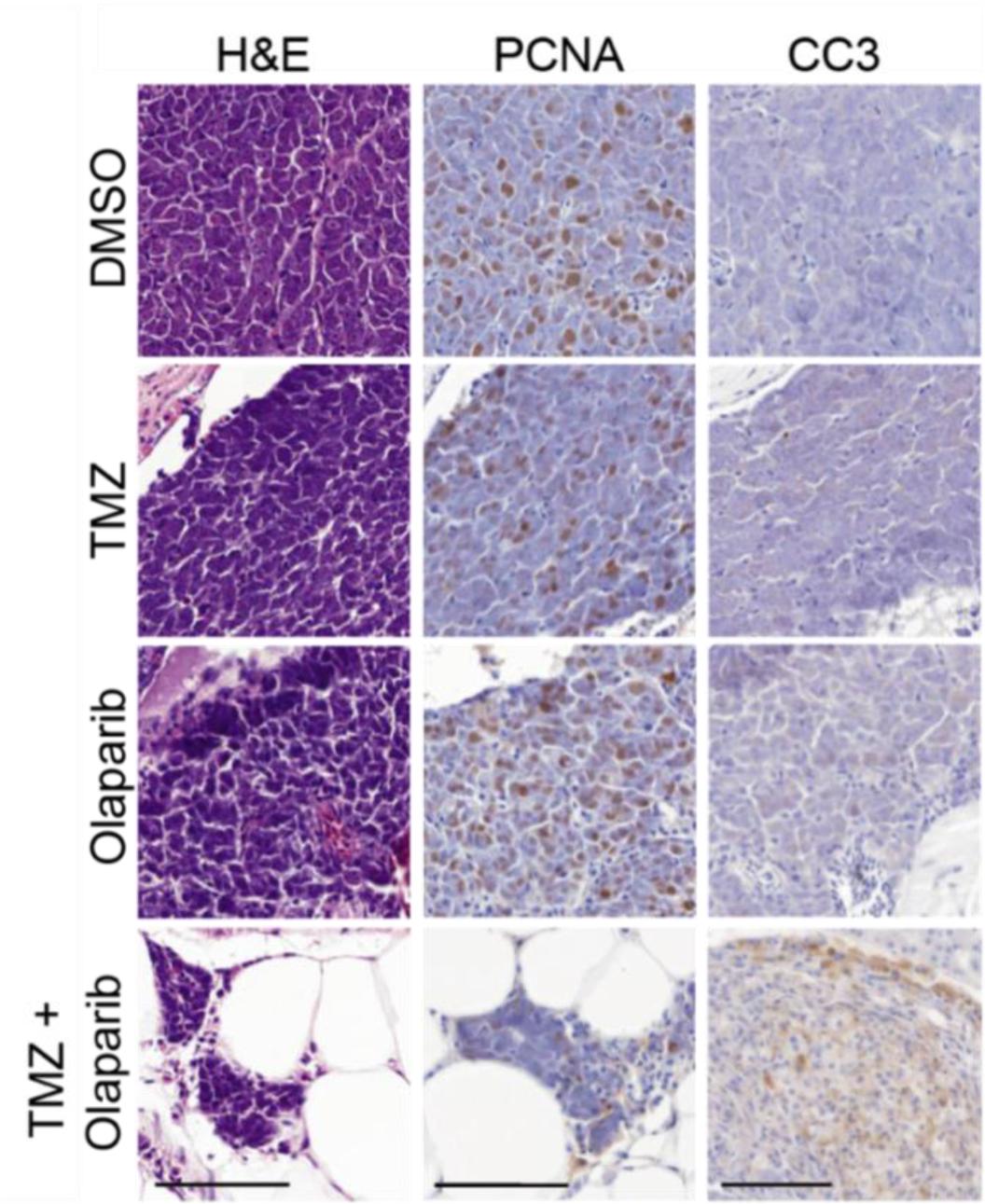
Hematoxylin and eosin (H&E), proliferating cell nuclear antigen (PCNA), and cleaved caspase 3 (CC3) staining of tumor sections from engrafted *EGFP;MYCN;tp53^-/-^;brca2^-/-^* NB tumor treated with vehicle control (DMSO), 33mg/kg TMZ, 50mg/kg olaparib, and 33mg/kg TMZ + 50mg/kg olaparib. Scale bars = 50μm.

**Supplementary Figure 6.**
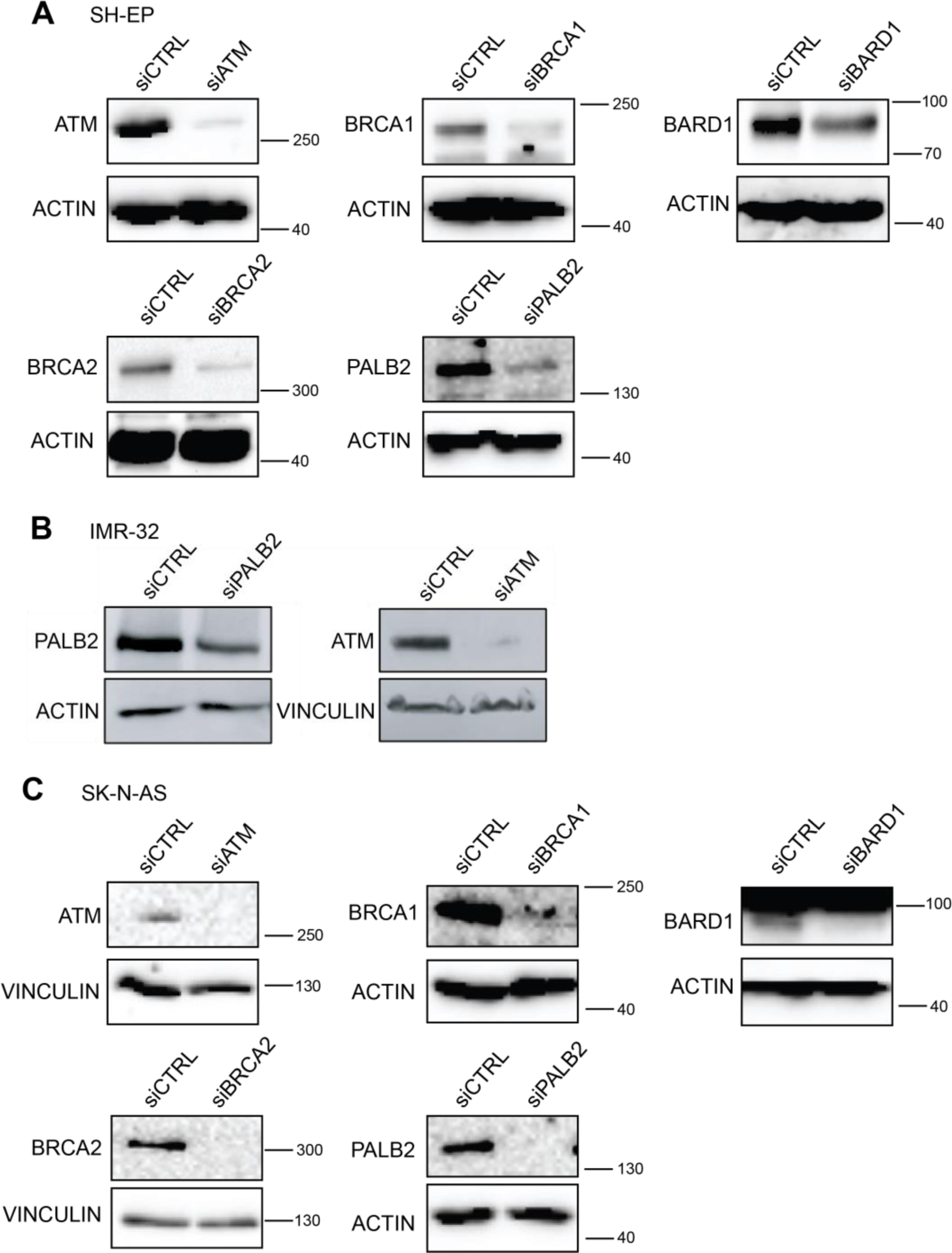
Confirmation of siRNA knock-down of DDR components (A) in SH-EP cells, (B) IMR-32 cells, and (C) SK-N-AS cells. siCTRL, non-targeting siRNA control.

**Supplementary Figure 7.**
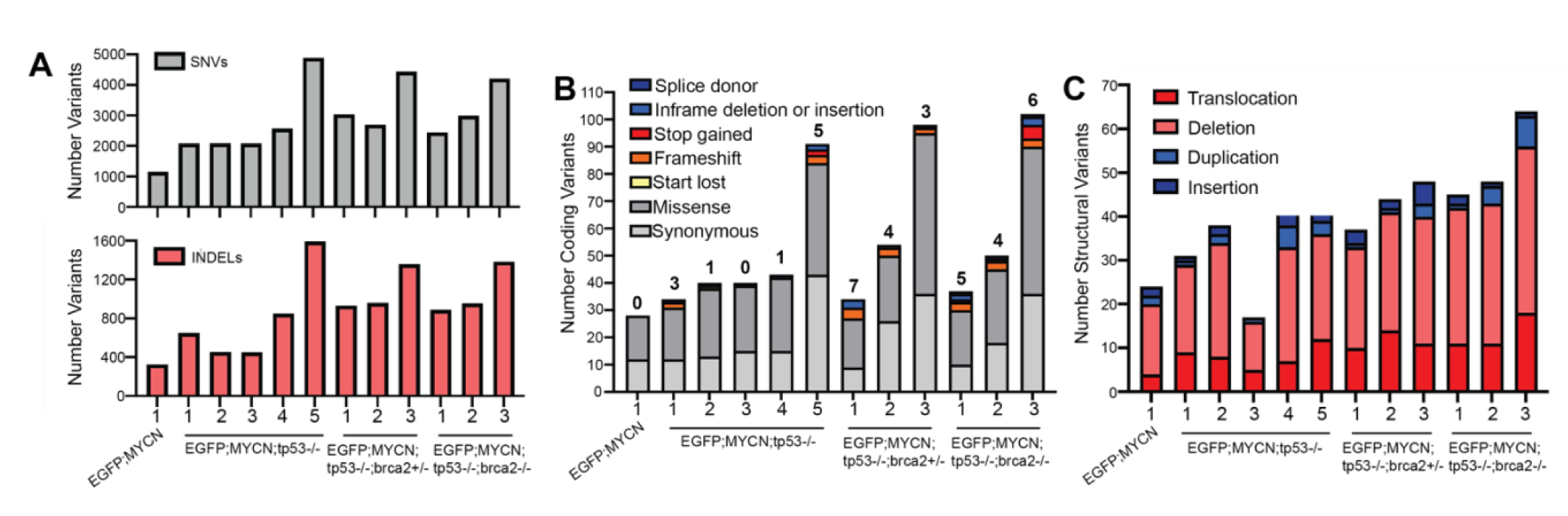
DDR-deficient zebrafish NB tumors display evidence of genomic instability. **(A)** Mutect2 identification and quantification of genome-wide single nucleotide variants (SNVs), and insertions and deletions (INDELS) for *EGFP;MYCN, EGFP;MYCN;tp53^-/-^; EGFP;MYCN;tp53^-/-^;brca2^+/-^,* and *EGFP;MYCN;tp53^-/-^;brca2^-/-^* tumors. **(B)** Mutect2 identification and quantification of coding variants for *EGFP;MYCN, EGFP;MYCN;tp53^-/-^; EGFP;MYCN;tp53^-/-^;brca2^+/-^,* and *EGFP;MYCN;tp53^-/-^;brca2^-/-^* tumors. Variant consequences are indicated, and quantification of total INDEL-type variants (inframe insertions or deletions, frameshift mutations) is numerically indicated above each bar, for each tumor. **(C)** Quantification of structural variants in *EGFP;MYCN, EGFP;MYCN;tp53^-/-^; EGFP;MYCN;tp53^-/-^;brca2^+/-^,* and *EGFP;MYCN;tp53^-/-^;brca2^-/-^* NBs.

**Supplementary Table 3.** Mutations in tumor cells of CRISPR/Cas9 mosaic-injected *EGFP;MYCN;tp53^-/-^* zebrafish. DNA was extracted from EGFP+ FACS sorted NB cells from *EGFP:MYCN:tp53^-/-^* fish injected with Cas9 protein and gRNAs targeting *brca2, atm, palb2,* or *bard1* at the one-cell stage. DNA fragments containing the target sequences were amplified and submitted for Sanger sequencing. Where multiple gRNAs were used spanning distant exons for *brca2* and *palb2*, each possible target site-specific INDELS as well as inter-exon deletions were tested using relevant 5’ and 3’ primer combinations (see Table S2 for gRNA and primer sequences), and shown for each tumor. Relative proportions of the total number of Sanger sequencing reads for each sequence is noted as a percentage, as estimated by ICE analysis (Synthego Performance Analysis, v3.0) and/or Indigo analysis (gear-genomics.com), where applicable.

| NB # | Tumor onset (wpf) | Gene | Target site | Mutation & Consequence |
| --- | --- | --- | --- | --- |
| 1 | 6 | *brca2* | 5' upstream | none, wild-type |
|  |  | *brca2* | exon 7 | none, wild-type |
|  |  | *brca2* | exon 10 | none, wild-type |
|  |  | *brca2* | Δ5'-exon7 | none detected |
|  |  | *brca2* | Δ5'-exon10 | none detected |
|  |  | *brca2* | Δexon7-10 | Deletion between exon 7 and 10 (*brca2*<sup>483_1081del</sup>), frame-shift and premature stop |
| 2 | 8 | *atm* | exon 54 | none, wild-type<br>4bp deletion (*atm*<sup>8176_8179del</sup>), frame-shift |
| 3 | 8 | *palb2* | exon 1 | none, wild-type |
|  |  | *palb2* | exon 5 | none, wild-type<br>Minor alleles/INDELS <5% each |
| | | *palb2* | Δexon1-5 | Deletion between exon 2 and 5 with<br>38bp insertion (*palb2*<sup>1_387delins38bp</sup>),<br>loss of START (ATG) codon, ~35%<br>Deletion between exon 2 and 5 with<br>3bp ACT insertion<br>(*palb2*<sup>6_359delinsACT</sup>), ~60% |
| 4 | 10 | *bard1* | exon 5 | 53bp deletion, 58bp insertion<br>(*bard1*<sup>1055_1107delins58bp*fs</sup>), frame-shift<br>and premature stop, ~70%<br>5bp insertion (*bard1*<sup>1101-</sup><br><sup>1102insAGAAC*fs</sup>), frame-shift and<br>premature stop, ~30% |

